# Ultrafast venous and sagittal sinus constrictions in the brain driven by abdominal pressure

**DOI:** 10.64898/2026.05.02.722426

**Authors:** Qingguang Zhang, Christopher S. Garborg, Noah Frank, Fatemeh Salehi Shahrbarbaki, Kevin L. Turner, Patrick J. Drew

## Abstract

Nearly all the blood supplying the cortex exits via the bridging veins (BVs) that drain into the superior sagittal sinus (SSS), making these vessels key chokepoints for cerebral blood flow. Using optical imaging in head-fixed mice, we found that the SSS and BVs exhibit ultrafast contractions (<0.1 s) at the onset of locomotion, following whisker stimulation, and upon awakening from sleep. Contractions of the BV and SSS were strongly correlated with abdominal muscle EMG activity and were tightly correlated with respiration at rest. The rapid decrease in blood volume caused by venous constrictions resulted in spurious increases in fluorescence in mice expressing fluorescent reporter proteins, creating artifacts that mimic functional signals. Venous contractions with the same amplitude and dynamics could be generated in anesthetized mice by abdominal pressure application, showing that these contractions were generated by mechanical coupling with the abdomen. Externally imposed abdominal pressures also drove a rapid but transient increase in blood flow. Unlike the pial and parenchymal microvasculature whose diameters are controlled by local signals, the diameters of SSS/BV are dynamically controlled during behavior by abdominal muscle regulation of intracranial pressure, establishing a pathway for regulation of cerebral hemodynamics via mechanical coupling between the central nervous system and the viscera.

## Introduction

Under physiological conditions, the flow of blood in the brain is regulated by signals from neurons and glia on a time scale of seconds to tens of seconds (1, 2). Relaxation of smooth muscle around arterioles, penetrating arteries and pial arteries is the most rapid of these processes, occurring within a few hundred milliseconds after neural activity increases (3–6). In behaving mice, pharmacological infusions silencing local neural activity block arterial dilation in the awake animal (7), showing arterial dilation is driven by local neural activity. Arterial dilation is followed by a much slower dilation of ascending and pial veins (4, 8) and capillaries (3, 9). Because of their sluggish (minutes) response to vasoconstrictors (10), the control of the cerebral venous system has received less attention and veins have been assumed to have slow and passive dynamics.

Blood flow from the dorsal cortex drains into the superior sagittal sinus (SSS), a large dural sinus formed by bifurcation of the dural membrane (**Figure 1**A). The SSS drains the dorsal and frontal portions of cortex (11) as well as dural vessels (12). The SSS is fed by the large bridging veins (BV) that run nearly perpendicular to the SSS. The bridging veins have sphincter-like rings of collagen surrounding the entrance of the bridging vein to the SSS (13, 14) which have been hypothesized to play a role in venous pressure regulation (15). Any changes in caliber of the SSS will change the vascular resistance and could thus change the blood flow for a large portion of cortex. The blood pressure decreases through the cerebral vascular network, so that in anesthetized animals (15–18) and in awake humans (19), SSS pressure is lower than intracranial pressure (ICP). This negative transmural pressure is maintained due to the relative stiffness of the venous wall which can resist collapse in the face of small negative pressure gradients, much like the hull of submarine allows the crew compartment to have a lower pressure than the water around it. Because nearly all the blood flowing through the cortex drains through the bridging veins and superior sagittal sinus, changes in the diameters of these vessels could have an influence on perfusion of all the cortex. In anesthetized animals, artificial elevation of ICP causes bridging veins to constrict (20). In behaving mice, ICP is dynamic and can rise from a baseline of 5-10 mmHg up to 25 mmHg or more during locomotion (21, 22). These ICP increases are driven by abdominal muscle contraction that drive increased intra-abdominal pressure which is transmitted to the central nervous system via a vascular network that spans the vertebral bones (23). Determining whether these behaviorally driven ICP increases have any impact on bridging veins and the SSS is important because diameter changes in these vessels could drive hemodynamic changes and affects fluid drainage.

**Figure 1.**
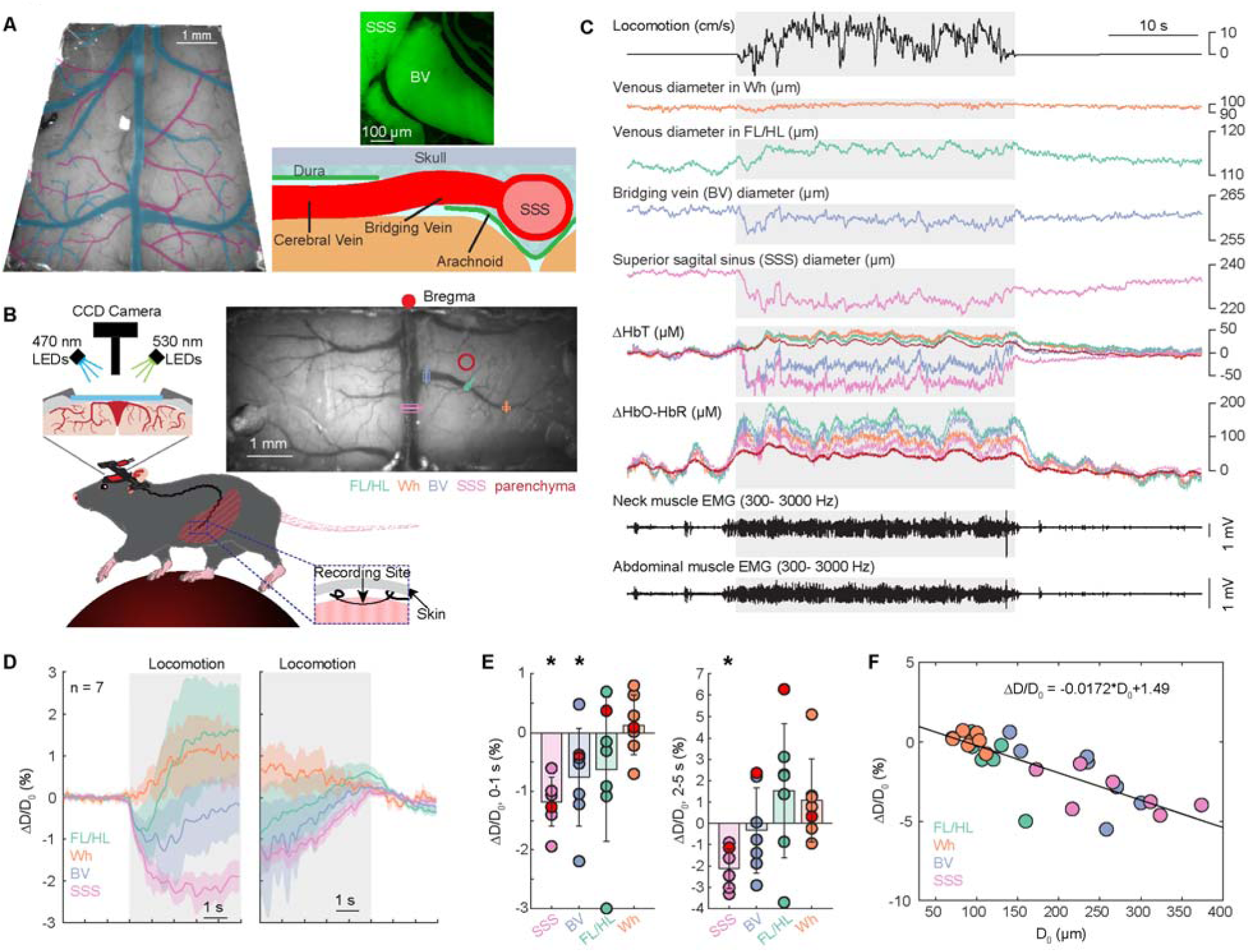
Venous constriction during voluntary locomotion in awake, head-fixed mice. (**A**) Schematic showing the pial vasculature in the mouse brain. Left, dorsal view *in vivo* under 530 nm illumination with veins traced in blue and arteries in magenta; Top right, images showing the dural venous sinuses after transcardial perfusion with FITC and gelatin and fixation; Bottom right, schematic showing the coronal view of the mouse brain. (**B**) Left, schematic of the experimental setup for widefield IOS imaging of awake head fixed mice. Right, image of the cerebral vasculature under 530 nm illumination through a thin-skull window spanning the parietal cortices of both hemispheres. Colored lines denote locations of vessel diameter measurements shown in subsequent figures. (**C**) Example data showing hemodynamic changes of veins at different locations (shown in **B**) during voluntary locomotion. FL/HL, forelimb/hindlimb representation of the somatosensory cortex; Wh, vibrissae cortex. ΔHbT, total hemoglobin; ΔHbO-HbR, differences of oxy- and deoxy-hemoglobin. The shaded area denotes the period of locomotion. (**D**) Averaged locomotion onset- and offset-triggered responses of ΔD/D_0_ (n = 7 mice) in veins at different locations. (**E**) Average venous diameter change (ΔD/D_0_) during the initial phase of locomotion (0-1 second after locomotion onset, left) and sustained locomotion (2-5 seconds after locomotion onset, right) in veins at different locations. There was a rapid constriction in superior sagittal sinus (SSS, -1.18 ± 0.42%, Wilcoxson rank sum test, p = 0.0003) and bridging veins (BV, -0.76 ± 0.84%, Wilcoxson rank sum test, p = 0.0084) in response to locomotion onset, while pial veins in the FL/HL (−0.63 ± 1.22%, Wilcoxson rank sum test, p = 0.09) and Wh (0.13 ± 0.51%, Wilcoxson rank sum test, p = 0.9327) did not change significantly. In response to constant locomotion, the initially constricted BV return to baseline level (−0.34 ± 2.01%, Wilcoxson rank sum test, p = 0.3450), while the SSS stayed constricted (−2.13 ± 0.92%, Wilcoxson rank sum test, p = 0.0003) and returned to baseline level within seconds after the cessation of voluntary locomotion. (**F**) Significant negative relationship (slope = -0.0172, 95% confidence interval [-0.0225, -0.0119], goodness of fit R^2^ = 0.6296, p = 4.7358e-7) between the venous diameter change and baseline vessel diameter, showing the large collecting vessels constrict in response to locomotion.

Here, we visualized the dynamics of bridging veins and the SSS using optical imaging in head fixed mice and found a constriction of the SSS and bridging veins (but not more lateral pial veins) with ultra-rapid onset (∼100ms) during bouts of voluntary locomotion. Sensory stimulation (to the whiskers) also drove abdominal muscle contraction and corresponding SSS/BV constriction, and in mice expressing fluorophores, this constriction decreased light absorbance by hemoglobin which could drive artifactual functional signals. The SSS/BV constriction was tightly correlated with abdominal muscle contraction (as measured with electromyography) and could be recapitulated in anesthetized mice by squeezing the abdomen. Our results show that, unlike the pial and parenchymal microvasculature whose diameters are controlled by local neural and glial signals, the diameters of SSS/BV are dynamically controlled during behavior by abdominal muscle regulation of intracranial pressure. Cerebral hemodynamics do not just reflect local neural activity, but also are impacted by abdominal muscle contractions via mechanical coupling between the central nervous system and the viscera.

## Results

We measured brain hemodynamics using widefield intrinsic optical imaging, two-photon laser scanning microscopy (2PLSM) and laser Doppler flowmetry in 7 head fixed C57BL/6 mice, as well as using mini-scope in one free-behaving C57BL/6 mouse. We also measured GFP responses in 6 head fixed CAG-EGFP mice. Intracranial pressure measurements in 10 C57BL/6 mice were from our previous publications (21, 22) and re-analyzed here.

### Voluntary locomotion drives rapid SSS/BV contraction and slower dilation of pial veins

We first assessed the spatial patterns of venous responses and their relationship to voluntary locomotion using widefield intrinsic optical signal (IOS) imaging. We performed imaging through thinned skull in awake C57BL/6 mice (n = 7 mice, 2 female) head fixed on a spherical treadmill (24, 25). Because the SSS is underneath the midline suture, the midline and coronal sutures were thinned to provide optical clarity. We illuminated the brain with alternating 530 nm and 470 nm light, providing information about blood volume, vessel diameter and brain oxygenation (24, 25). We calculated the vessel diameter changes using the full-width at half maximum (FWHM) (25, 26) from the IOS images under 530 nm illumination. We recorded EMG signals from the abdominal musculature, which are engaged during locomotion and drive brain motion (23).

We first looked at the vascular responses to voluntary bouts of locomotion, averaging vascular changes evoked by movement into locomotion triggered averages. Consistent with our previous results (8, 21, 27), voluntary locomotion evokes a large increases in arterial diameter within a few hundred milliseconds (**Supplementary Figure 1**A). Both the abdominal muscles (23) and nuchal muscle showed a strong increase in activity during locomotion (**Figure 1**C and **Supplementary Figure 4**B). In contrast, there were branch-specific changes in venous diameter (ΔD/D_0_) in response to locomotion (**Figure 1**C-E). Specifically, the smaller more lateral veins (in the whisker and forelimb/hindlimb representations) dilated during locomotion, while the SSS and bridging veins rapidly constricted. At the cessation of locomotion, all the veins returned to their baseline diameter (**Figure 1**D). The constriction of SSS and BV occurred prior to any vasodilation in the somatosensory cortex (**Figure 1**C and D). Similar constrictions were observed using two-photon microscopy (**Supplementary Figure 1**A-C). Moreover, we also observed bridging vein constriction in a freely moving mouse during locomotion using a miniscope (**Supplementary Figure 1**E), showing the observed responses are not due to body posture difference caused by head fixation. As an alternate metric of vasodilation, we quantified the change in blood volume using changes of total hemoglobin (ΔHbT, **Supplementary Figure 2**) around these veins, and saw similar dynamics, with the decrease in blood volume in SSS and BV, and increase in the smaller, more lateral veins (**Supplementary Figure 2**B, C and E). Interestingly, though the blood volume decreased in both the SSS and BV, the blood oxygenation (ΔHbO-HbR) increased (**Supplementary Figure 2**D and F). The contractions of these veins were ultrarapid (∼100 ms), far too quickly to be due to mural cell contractions (3, 9), so we sought other mechanisms. We observed a significant negative correlation between the basal vein diameter and its diameter change during locomotion (**Figure 1**F). Because the smaller veins feed into larger veins, the blood pressure inside the vein decreases with size, while the pressure outside all the veins (the intracranial pressure) will be roughly constant. If the ICP rises enough, then the balance of pressures favors a decrease in the diameter in the larger veins. Movements in both humans and mice are often preceded by the activation of abdominal muscles (28), whose contraction stiffens the core in anticipation of movement. Previous studies have shown that intraabdominal pressure (IAP) is coupled to ICP (29) and abdominal muscle contraction drives increases in IAP (28), leading to forces on the brain that drive brain motion (23) and increase ICP (30). To understand how these muscle contractions were related to the rapid changes in venous diameter, we recorded oblique abdominal muscle electromyograph (EMG_Abd_) signals (n = 4 mice, all male) during locomotion. We observed a robust increase in EMG_Abd_ power that are strongly correlated with locomotion (**Figure 1**C and **Supplementary Figure 4**B). When we compare the cross-correlation curve between venous constriction and abdominal EMG activity (**Supplementary Figure 4**D) or locomotion speed (**Supplementary Figure 4**E), we observed that the constriction of SSS invariably followed EMG activity, but often preceded locomotion, suggesting that abdominal muscle contraction drives the SSS constriction. A network of veins connects the abdominal cavity to the spinal column, functioning as a hydraulic system that can transfer force between the two compartments (23). Abdominal muscle contraction increases abdominal pressure applies force (31) to the abdomen (which drives a pressure increase) which is transmitted to the central nervous system by a network of veins connecting the abdominal cavity to the spinal column, functioning as a hydraulic system that can transfer force between the two components (23). Our results suggest that this abdominally driven force also drives contraction of the large veins in the cortex.

### Whisker stimulation drives contraction of BV and SSS

Sensory evoked responses can drive both covert and overt muscle contractions (32, 33). Since many imaging experiments make use of sensory-evoked responses, such as whisker stimulation (34), we then asked if constriction of the SSS and BV could be driven by sensory stimulation of the whiskers, potentially mediated by abdominal muscle contractions. Brief (0.1s at 10 psi) unilateral stimulation of the whiskers drive vasodilation of arteries in the contralateral sensory cortex with a ∼1 second delay (4). However, we also observed a rapid (< 0.2 seconds of stimulation onset) constriction in the SSS and BV, but not of the smaller veins in the whisker representation of the somatosensory cortex (**Figure 2**B and C). Quantification of ΔHbT also showed a decrease in response around stimulation onset (**Supplementary Figure 3**B, C and E), and in contrast to locomotion, there was a brief dip in net oxygenation of the blood (**Supplementary Figure 3**D and F), likely because the stimulus was unexpected and was not accompanied by an increase in respiration that raises systemic blood oxygenation (25). The dip in oxygenation is reminiscent of the “initial dip” seen in some imaging experiments (35). Electromyography recordings showed a large increase in abdominal EMG (EMG_Abd_) activity time locked to the whisker stimulation (**Figure 2**B and **Supplementary Figure 4**C). As a control we looked at the responses to auditory stimulation, which drives whisking and similar (though somewhat smaller) functional hyperemia in the whisker representation of the cortex due to whisking (4, 36). Auditory stimulation did not drive abdominal EMG activation (**Figure 2**B and **Supplementary Figure 4**C), and there was no constriction of the SSS and BV (**Figure 2**B), consistent with the hypothesis that abdominal muscle constriction drove the constriction of the largest veins.

**Figure 2.**
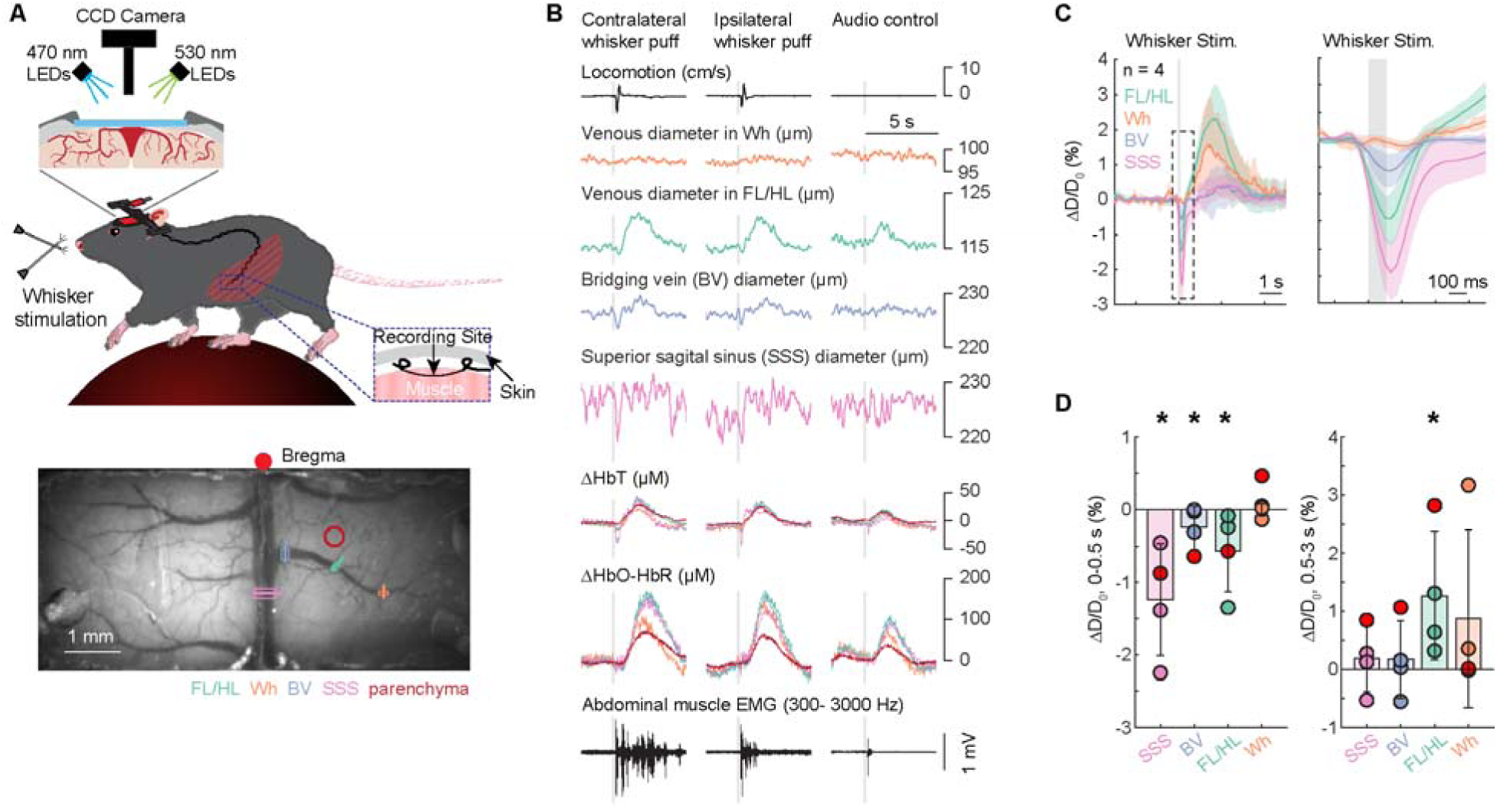
Sagittal sinus and bridging vein constriction in response to whisker stimulation in awake, head-fixed mice. (**A**) Top, schematic of the experimental setup for widefield IOS imaging of awake head fixed mice during whisker stimulation. Bottom, image of the cerebral vasculature under 530 nm illumination through a thin-skull window spanning the parietal cortices of both hemispheres. Colored lines denote locations of vessel diameter measurements shown in subsequent figures. (**B**) Example single trial responses showing hemodynamic changes of veins at different locations (shown in **A**) in response to whisker stimulation. FL/HL, forelimb/hindlimb representation of the somatosensory cortex; Wh, vibrissae cortex. ΔHbT, total hemoglobin; ΔHbO-HbR, differences between oxy- and deoxy-hemoglobin. The shaded area denotes the period of whisker stimulation. (**C**) Group average of a short (100 ms) whisker stimulation triggered response of ΔD/D_0_ (n = 4 mice) in veins at different locations. Inset showing a zoom-in view of ΔD/D_0_ responses immediately before (330 ms) and after (660 ms) whisker stimulation. (**D**) Average venous diameter change (ΔD/D_0_) during the initial phase of whisker stimulation (0-0.5 second after locomotion onset, left) and after stimulation (0.5-3 seconds after whisker stimulation onset, right) in veins at different locations. There was a rapid constriction in superior sagittal sinus (SSS, -1.24 ± 0.77%, Wilcoxson rank sum test, p = 0.0143), bridging veins (BV, - 0.25 ± 0.30%, Wilcoxson rank sum test, p = 0.0143) and pial veins in the FL/HL (−0.56 ± 0.56%, Wilcoxson rank sum test, p = 0.0143) in response to a short whisker stimulation, while pial veins in the Wh (−0.005 ± 0.09%, Wilcoxson rank sum test, p = 0.9429) did not change significantly. After the initial constriction, SSS (0.18 ± 0.57%, Wilcoxson rank sum test, p =0.1571) and BV (0.17 ± 0.67%, Wilcoxson rank sum test, p =0.1571) return back to baseline levels, while the pial veins in FL/HL (1.27 ± 1.11%, Wilcoxson rank sum test, p = 0.0143) dilated. The pial veins in the Wh did not change significantly (0.88 ± 1.54%, Wilcoxson rank sum test, p = 0.1571).

### SSS and BV constrictions can generate artefactual fluorescence increases with kinetics similar to functional indicators

Hemoglobin absorbs light, and changes in hemoglobin levels will change the absorbance of excitation and emission light (37), which may confound interpretation of neural activity using fluorescent calcium indicators (38–41). This is particularly salient for the SSS/BV constrictions we observe, which have onsets and offset similar to genetically-encoded calcium indicators (42), rather than the slower dynamics of blood volume changes (27). To investigate the role of venous constriction in generating fluorescence artifacts, we quantified the effects of hemodynamics on fluorescence measured from CAG-EGFP mice (n = 6, all male) with widefield fluorescence imaging in response to whisker stimulation head fixed in a tube (**Figure 3**A). Note that in this preparation, the whisker puff evoked abdominal EMG responses (**Supplementary Figure 5**B) is slightly smaller than these mice head fixed on a spherical treadmill (**Supplementary Figure 4**C). Because green fluorescence protein (GFP) is not responsive to neuronal activity, changes in fluorescence will be entirely due to changes in the local concentration of light-absorbing hemoglobin. We observed large amplitude fluctuations that were restricted mainly to the midline vasculature and typically included fast transient spikes in fluorescence (**Figure 3**B). These fluctuations resulted from changes in venous blood volume along the SSS that relate to movements or postural changes (**Figure 3**B-D). We then tested three commonly used methods for correcting hemodynamics contamination: (1) hemodynamic correction using only green reflectance (i.e., single-wavelength method)(43), (2) hemodynamic correction using estimated excitation and emission attenuation (i.e., Ex-Em method)(40, 41), and (3) hemodynamic correction using a spatially detailed regression-based method to estimate hemodynamics contamination (i.e., spatial model)(38). The spatial model is based on a linear form of the Beer-Lambert relationship but is fit at every pixel in the image and does not rely on the estimation of physical parameters. For the artifacts caused by initial rapid constrictions of veins and slow dilation of veins, both the spatial model and regression model can successfully remove the artifact, but the Beer-Lambert method can cause over correction (**Figure 3**B-F).

**Figure 3.**
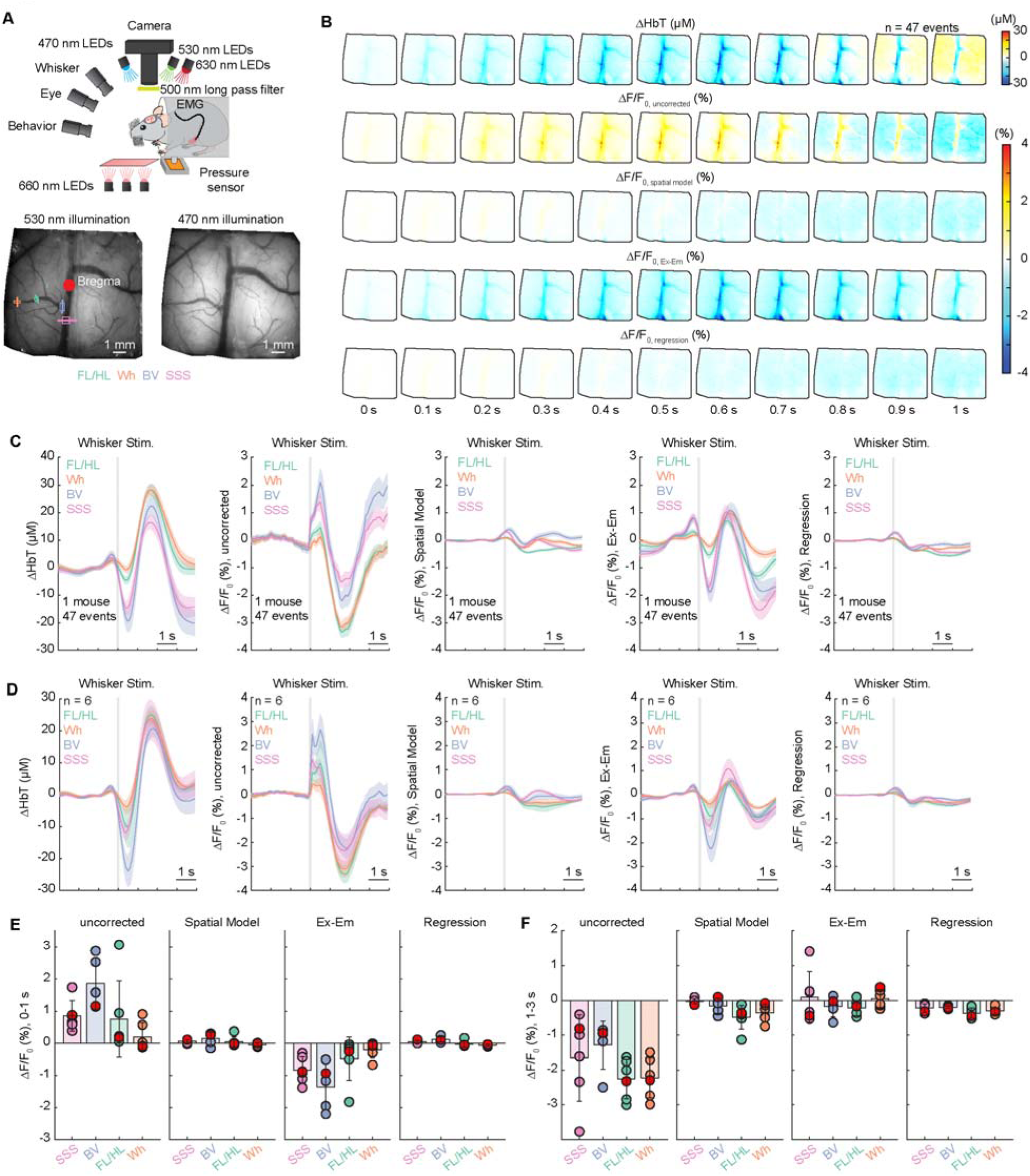
Ultrafast venous constriction induces contamination in widefield calcium imaging. (**A**) Top, schematic of the experimental setup for widefield IOS imaging of awake head fixed mice during whisker stimulation. Bottom, image of the cerebral vasculature under 530 nm illumination (bottom left) and image of the fluorescent signals under 470 nm illumination (bottom right) through a thin-skull window spanning the cortices of both hemispheres. Colored lines denote locations of vessel diameter measurements shown in subsequent figures. (**B**) Averaged spatial distribution of ΔHbT, raw fluorescence signal (ΔF/F_0, uncorrected_) and corrected fluorescence signal using different methods (ΔF/F_0, SpatialModel_, ΔF/F_0, Ex-Em_ and ΔF/F_0, Regression_) in response to contralateral whisker stimulation (n = 47 events in 1 mouse). (**C**) An example showing averaged trace of contralateral whisker stimulation triggered response of ΔHbT, raw fluorescence signal and corrected fluorescence signal using different methods (n = 47 events in 1 mouse) in veins at different locations in the same animal shown in (**B**). (**D**) Group (n = 6 mice) average of whisker stimulation triggered response of ΔHbT, raw fluorescence signal and corrected fluorescence signal using different methods in veins at different locations. (**E**) Average fluorescence change (ΔF/F_0_) during the initial phase of contralateral whisker stimulation (0-0.8 second after whisker stimulation onset) in veins at different locations. During the initial phase after the whisker stimulation, we observed a fast increase of fluorescence in superior sagittal sinus (SSS, 0.85 ± 0.47%, Wilcoxson rank sum test, p = 0.0011), bridging veins (BV, 1.86 ± 0.81%, Wilcoxson rank sum test, p = 0.0022) and in pial veins in the FL/HL (0.75 ± 1.19%, Wilcoxson rank sum test, p = 0.0011), while fluorescence of and pial veins in the Wh (0.20 ± 0.43%, Wilcoxson rank sum test, p = 0.5498) did not change significantly. These artifactual fluorescence signals can be corrected using spatial model method (SSS, 0.05 ± 0.05%, paired t test, p = 0.0085; BV, 0.15 ± 0.20%, paired t test, p = 0.0133) and regression method (SSS, 0.04 ± 0.05%, paired t test, p = 0.0080; BV, 0.11 ± 0.08%, paired t test, p = 0.0068). However, the Ex-Em method over-corrected these signals (SSS, -0.84 ± 0.41%, paired t test, p = 0.0045; BV, -1.35 ± 0.71%, paired t test, p = 0.0076). (**F**) Average fluorescence change (ΔF/F_0_) during 1-3 seconds after whisker stimulation in veins at different locations. During later phase after the whisker stimulation, we observed a decrease of fluorescence signals in superior sagittal sinus (SSS, -1.65 ± 1.24%, Wilcoxson rank sum test, p = 0.0011), bridging veins (BV, -1.28 ± 0.69%, Wilcoxson rank sum test, p = 0.0022), pial veins in the FL/HL (−2.26 ± 0.57%, Wilcoxson rank sum test, p = 0.0011), and pial veins in the Wh (−2.23 ± 0.58%, Wilcoxson rank sum test, p = 0.0011). These artifactual fluorescence signals can be corrected using spatial model method (SSS, -0.02 ± 0.08%, paired t test, p = 0.0232; BV, -0.15 ± 0.22%, paired t test, p = 0.0092; FL/HL, -0.49 ± 0.34%, paired t test, p = 0.0001; Wh, -0.36 ± 0.25%, paired t test, p = 0.0002), regression method (SSS, -0.22 ± 0.12%, paired t test, p = 0.0380; BV, -0.20 ± 0.04%, paired t test, p = 0.0209; FL/HL, -0.37 ± 0.15%, paired t test, p = 0.0001; Wh, -0.30 ± 0.10%, paired t test, p = 0.0002;), and the Ex-Em method (SSS, 0.10 ± 0.73%, paired t test, p = 0.0750; BV, - 0.18 ± 0.29%, paired t test, p = 0.0374; FL/HL, -0.22 ± 0.22%, paired t test, p = 0.0001; Wh, 0.05 ± 0.25%, paired t test, p = 0.0003).

### Intracranial pressure dynamics during behavior

The rapid contraction of veins could be mediated by increases in intracranial pressure (ICP), as artificial elevations of ICP are known to compress bridging veins (20) and ICP is elevated during locomotion (21, 22, 24). To address this, we reanalyzed a previously published dataset (21, 22) of intracranial pressure recordings made from mice head fixed on a spherical treadmill. In the reanalysis, we used broadband signals with minimal temporal filtering in order to capture the fast temporal dynamics. An example ICP trace, along with locomotion dynamics was shown in **Figure 4**A. It is apparent that there are large (>10 mmHg) increases in ICP during locomotion that dominate all other fluctuations, consistent with ICP increases being the drives of the contractions of the largest veins. The locomotion triggered averages show an increase in ICP that precedes the onset of locomotion, rises ∼8 mmHg above the resting pressure, and then returns to baseline shortly after the cessation of locomotion (**Figure 4**C), consistent with the dynamics of the venous contractions during locomotion and EMG activation. Inspection of periods of quiescence and of the power spectrum reveal ∼3 Hz fluctuations in ICP that are less than 1 mmHg (0.41 ± 0.11 mmHg, n = 10 mice) in amplitude (**Figure 4**A and B). There are also ∼10 Hz fluctuations (at the cardiac frequency) visible in the power spectrum (**Figure 4**B) that are at least an order of magnitude smaller than the respiration fluctuations, too small to have an appreciable effect on venous diameter. While both cardiac and respiratory-induced pressure fluctuations are at least an order of magnitude smaller than those that accompany locomotion, the ∼1mmHg fluctuations due to respiration could potentially have an impact on venous volume (44, 45). To address this, we measured respiration with a thermocouple (24, 25) during widefield imaging. Increases and decreases in the thermocouple temperature correspond to the expiratory and inspiratory phases of the respiratory cycle, respectively (**Figure 4**D). During resting period, there was substantial correlation between the diameter of the SSS, BV and pial veins in FL/HL, and the respiration signal (**Figure 4**D and E). Intra-abdominal pressure (and therefore ICP) rises during inspiration (46), which tends to compress the SSS, though these pressure fluctuations are much smaller than those caused by locomotion. We observed that the diameter of the cerebral veins (especially SSS and BV) depends on respiratory cycles (it decreases during inspiration, and increases during expiration). These results show that the drivers of ICP increase (locomotion and respiration) are correlated with BV/SSS dynamics.

**Figure 4.**
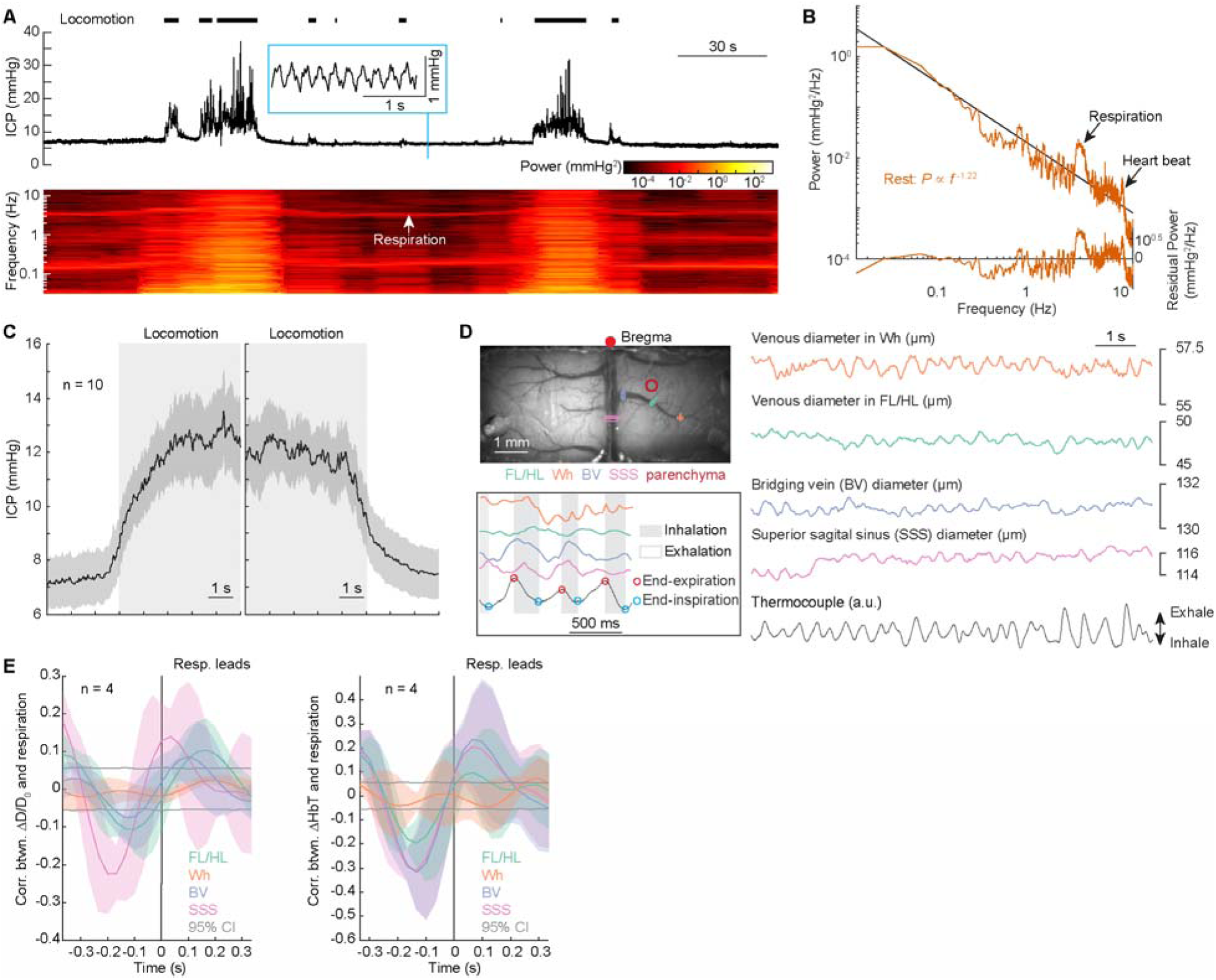
Intracranial pressure increases during locomotion and fluctuates with respiration. (**A**) Example intracranial pressure (ICP) dynamics during voluntary locomotion. Top, black trace shows ICP, and black tick marks show locomotion events. Inset showing a zoom-in of respiration-driven ICP oscillations. Bottom, spectrogram of ICP during rest and locomotion. (**B**) Representative power spectrum of the ICP as well as its power law fit (dashed line) using the data from resting period. The residual power (i.e., the difference between the power spectrum of the observed ICP signal and the power law fit) is shown in the bottom. (**C**) Locomotion onset and offset triggered ICP changes. Note that ICP rises before locomotion onset (onset time: -0.56 ± 0.29 s). (**D**) Example trace showing the diameter change of veins at different locations is correlated with respiration. Inset, a zoom in view of the temporal relationship between respiration and venous diameters. The respiration signal here is from the temperature change measured outside the nostril, i.e., an increasing signal is the expiration phase, and decreases in the temperature happen during the inspiration phase. (**E**) Left, correlation coefficient between respiration signal and venous diameter change. Right, correlation coefficient between respiration signal and venous blood volume (ΔHbT). A positive peak with positive time lag suggests that upwarding respiration signal (i.e., expiration) causes venous dilation, and downwarding respiration signal (i.e., inspiration) causes venous constriction.

### SSS and bridging veins dilate during sleep and constrict upon awakening

Next, we looked the dynamics of the large veins during sleep. Sleep is accompanied by large dilations of arteries (47, 48) and global increases in blood volume (49) that are much larger than what are seen in the awake state. Sleep, particularly rapid eye movement (REM) sleep, is accompanied by large decreases in muscle tone (50), and drops in blood pressure. We hypothesized that the net effect of these changes during sleep would drive dilation of the SSS and BV. We performed widefield imaging on head-fixed mice (n = 4, all male) in a tube (36, 47) and tracked whisker position, body movement, and abdominal muscle EMG (**Figure 5**A), which were used to determine the arousal state of the animal. We performed sleep scoring as previously described (47, 48) and categorized the arousal state into three stages: awake, rapid eye movement (REM) and non-rapid eye movement (NREM) sleep. In contrast to voluntary locomotion and whisker stimulation evoked venous constrictions, we found venous dilation associated with sleep, and saw large constrictions upon transitions from REM or NREM to the awake state (**Figure 5**B-D). We also saw large decreases in total hemoglobin and, in contrast to the constrictions during locomotion, decreases in blood oxygenation around these vessels in transitions to the awake state (**Figure 5**B-D). As with the venous dynamics in the awake state, constrictions were associated with higher abdominal EMG power (**Figure 5**B and C), again suggesting that there is abdominal control of venous dynamics across many arousal states.

**Figure 5.**
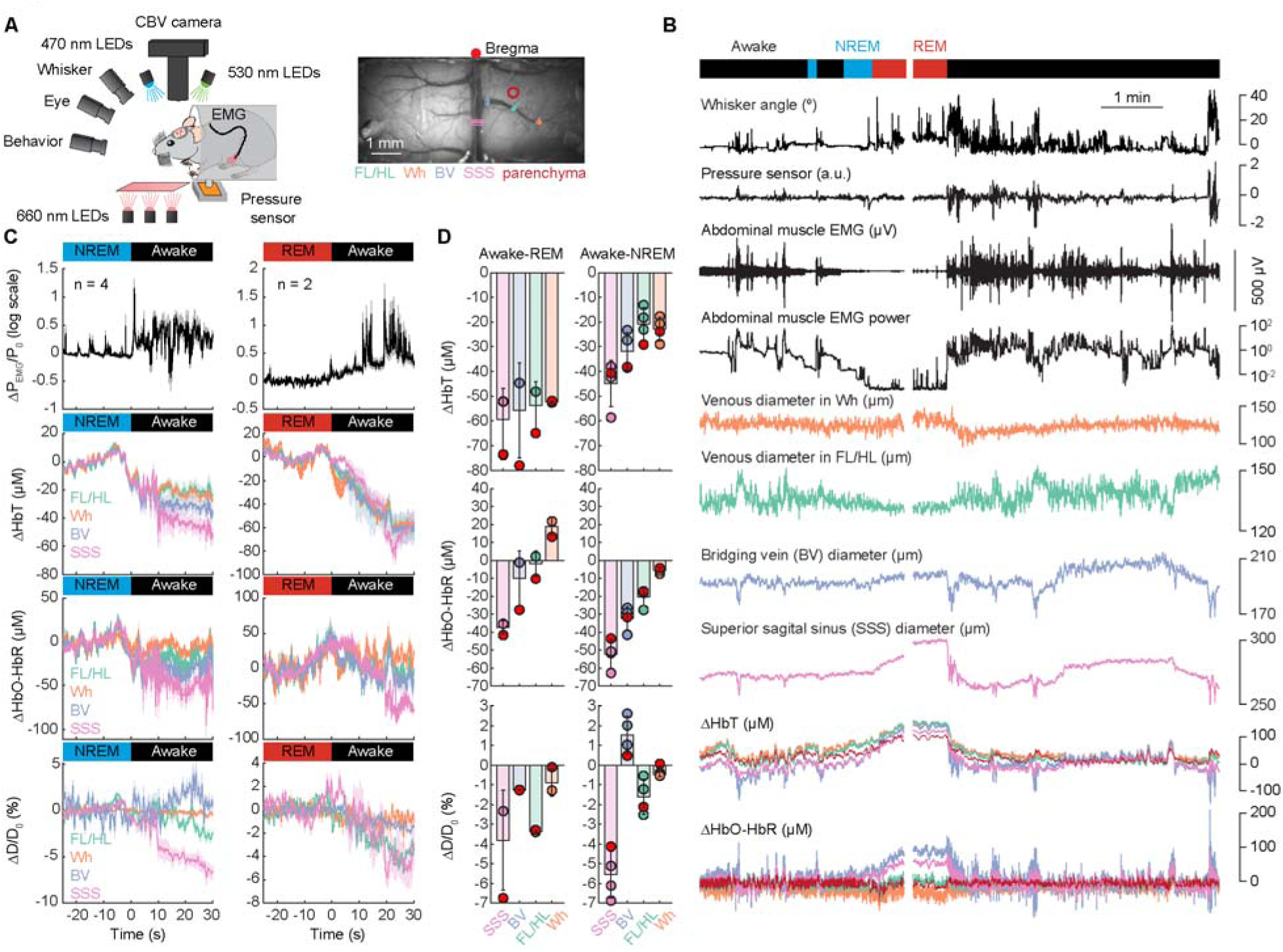
Awakening from NREM and REM sleep drives venous constriction. (**A**) Left, schematic of the experimental setup for widefield IOS imaging of head fixed mice. Right, image of the cerebral vasculature under 530 nm illumination through a thin-skull window spanning the parietal cortices of both hemispheres. Colored lines denote locations of vessel diameter measurements shown in subsequent figures. (**B**) Example data showing hemodynamic changes of veins at different locations (shown in **A**) during transitions between arousal states. Top, arousal sate scored from EMG/ECoG/whisker and body motion. White break denotes break in data collection between trials. FL/HL, forelimb/hindlimb representation of the somatosensory cortex; Wh, vibrissae cortex. ΔHbT, total hemoglobin; ΔHbO-HbR, differences of oxy- and deoxy-hemoglobin. The shaded area denotes the periods of different arousal states. (**C**) Changes of abdominal muscle EMG (ΔP_EMG_/P_0_), ΔHbT, ΔHbO-HbR, ΔD/D_0_ during the transition of awake into NREM (n = 4 mice), NREM into REM (n = 2 mice), NREM into awake (n = 4 mice), REM into awake (n = 2 mice). Note that scales are different across conditions. (**D**) Bar plot showing the mean changes in diameter and hemodynamic signals in response to each arousal state transition.

### Passively applied abdominal pressure drives SSS and BV constriction and increases in blood flow

Externally applied abdominal pressure drives brain movement like that seen during behavior, mediated through a network of valveless veins (vertebral venous plexus) that link the spinal cavity with the abdominal cavity (23). We asked if this pressure application could drive contraction of the SSS and BV. We anesthetized mice (1% isoflurane) and placed them in a computer controlled pneumatic pressure cuff (23) and applied one-second long squeezes of the abdomen while monitoring both cortical vessel diameter and cerebral blood flow (CBF) using laser Doppler flowmetry (51, 52). During cuff inflation (1 s), the pressure applied to the abdominal muscle increases and reaches half-max level within ∼0.17s. We found that externally applied abdominal pressure drove reliable constrictions of the SSS and BV, but not in the smaller pial veins (**Figure 6**B-D), recapitulating the patterns seen during locomotion (**Figure 1** and **Supplementary Figure 2**) and during sensory stimulation (**Figure 2** and **Supplementary Figure 3**). Interestingly, the constriction also drove a flow increase in the parenchyma (3.02 ± 1.47%, **Figure 6**F) and oxygenation increases across the whole cortex (**Figure 6**E). It seems that the abdominal pressure can drive a very rapid blood flow increase, and this could be a mechanism by which flow to the brain is increased peremptorily by abdominal muscle contractions, such as those that accompany the startle response (53).

**Figure 6.**
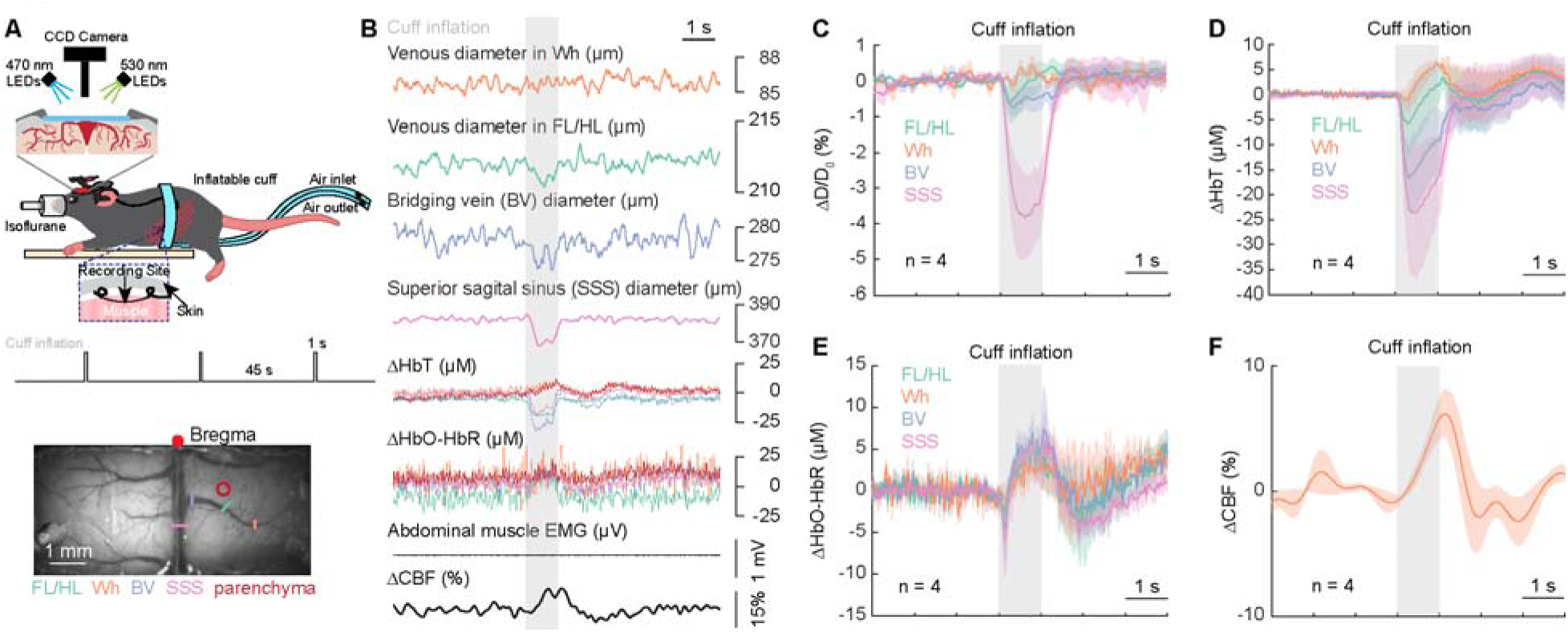
Abdominal squeezing cause SSS constrictions. (**A**) Top, schematic of the experimental setup for abdominal compression experiments. Bottom, image of the cerebral vasculature under 530 nm illumination through a thin-skull window spanning the parietal cortices of both hemispheres. Colored lines denote locations of vessel diameter measurements shown in subsequent figures. (**B**) Example trace showing brain hemodynamics changes at different locations during abdominal compression. (**C**) Population average of abdominal compression evoked responses of ΔD/D_0_ (n = 4 mice). Brief, gentle compression of abdomen reduced venous diameter in SSS (−3.18 ± 1.89%, Wilcoxson rank sum test, p = 0.0143) and BV (−0.53 ± 0.51%, Wilcoxson rank sum test, p = 0.0143), and no change of veins in FL/HL (−0.10 ± 0.79%, Wilcoxson rank sum test, p = 0.9429) and Wh (0.19 ± 0.04%, Wilcoxson rank sum test, p = 1). (**D**) As in (**C**) but for ΔHbT. Brief, gentle compression of abdomen reduced cerebral blood volume in SSS (−18.98 ± 19.08 µM, Wilcoxson rank sum test, p = 0.0143) and BV (−11.63 ± 11.31 µM, Wilcoxson rank sum test, p = 0.0143), but no change of veins in FL/HL (−1.06 ± 8.59 µM, Wilcoxson rank sum test, p = 0.9429) and Wh (3.27 ± 2.64 µM, Wilcoxson rank sum test, p = 0.9429). (**E**) As in (**C**) but for ΔHbO-HbR. Brief, gentle compression of abdomen did not change brain oxygenation in SSS (3.41 ± 3.15 µM, Wilcoxson rank sum test, p = 0.1571), but increased brain oxygenation in BV (4.55 ± 3.09 µM, Wilcoxson rank sum test, p = 0.0143), in veins in FL/HL (3.09 ± 0.91 µM, Wilcoxson rank sum test, p = 0.0143) and Wh (2.01 ± 1.40 µM, Wilcoxson rank sum test, p = 0.0143). (**F**) As in (**C**) but for ΔCBF. Significant increases of ΔCBF were observed in the parenchyma in response to 1 s squeezing (0.2-1.2 seconds after squeezing onset, 3.02 ± 1.47%, Wilcoxson rank sum test, p = 0.0143).

## Discussion

We observed rapid constrictions of bridging veins and the superior sagittal sinus during voluntary locomotion, whisker stimulation, and upon awakening from sleep, all conditions were accompanied by abdominal muscle contraction. Externally imposed abdominal pressure generated patterns of venous constriction that were very similar in amplitude compared to those seen during locomotion and sensory stimulation. These results are consistent with a model where abdominal pressures are transmitted to the central nervous system. These forces can cause brain movement (23), and here we show they can cause compression of the largest veins on a sub-second time scale.

Previous studies looking at more lateral and smaller veins have observed dilation (4, 51), something we have seen here as well. The cause of the size dependent difference in venous responses is likely due to differences in blood pressure throughout the network. Blood pressure decreases monotonically as blood transits through the vascular network (54), so the larger veins will have a lower pressure than the smaller veins that feed them. This means that if ICP is elevated, then first vessels in the brain to be compressed will be the large veins, as was observed here. ICP changes will depend on the coupling from the abdominal cavity to the spinal cord (**Figure 7**A). These results, coupled with the observed ICP increases during locomotion, point to a model where the central nervous system is mechanistically linked to the abdominal cavity, and pressures generated in the abdomen are transmitted to the brain, leading to increase in ICP and brain motion (23). If the ICP rises, it will be above the intraluminal pressures of the bridging veins and SSS (but not above the intraluminal pressures of the smaller veins)(**Figure 7**B, C and D), causing a spatial pattern of venous responses like the one we see during locomotion (**Figure 1**F). Notably, there are several caveats to our work. The superior sagittal sinus does not have a circular cross section, it is roughly triangular in histological sections, so our estimates of diameter changes may over or underestimate the actual changes. There are other large veins in the brain that we did not image (e.g., transverse sinuses) that may undergo similar ICP-driven compression. Finally, ICP will not be completely spatially homogenous, and the venous pressure could vary dynamically, which will affect the sensitivity of veins to elevations in ICP.

**Figure 7.**
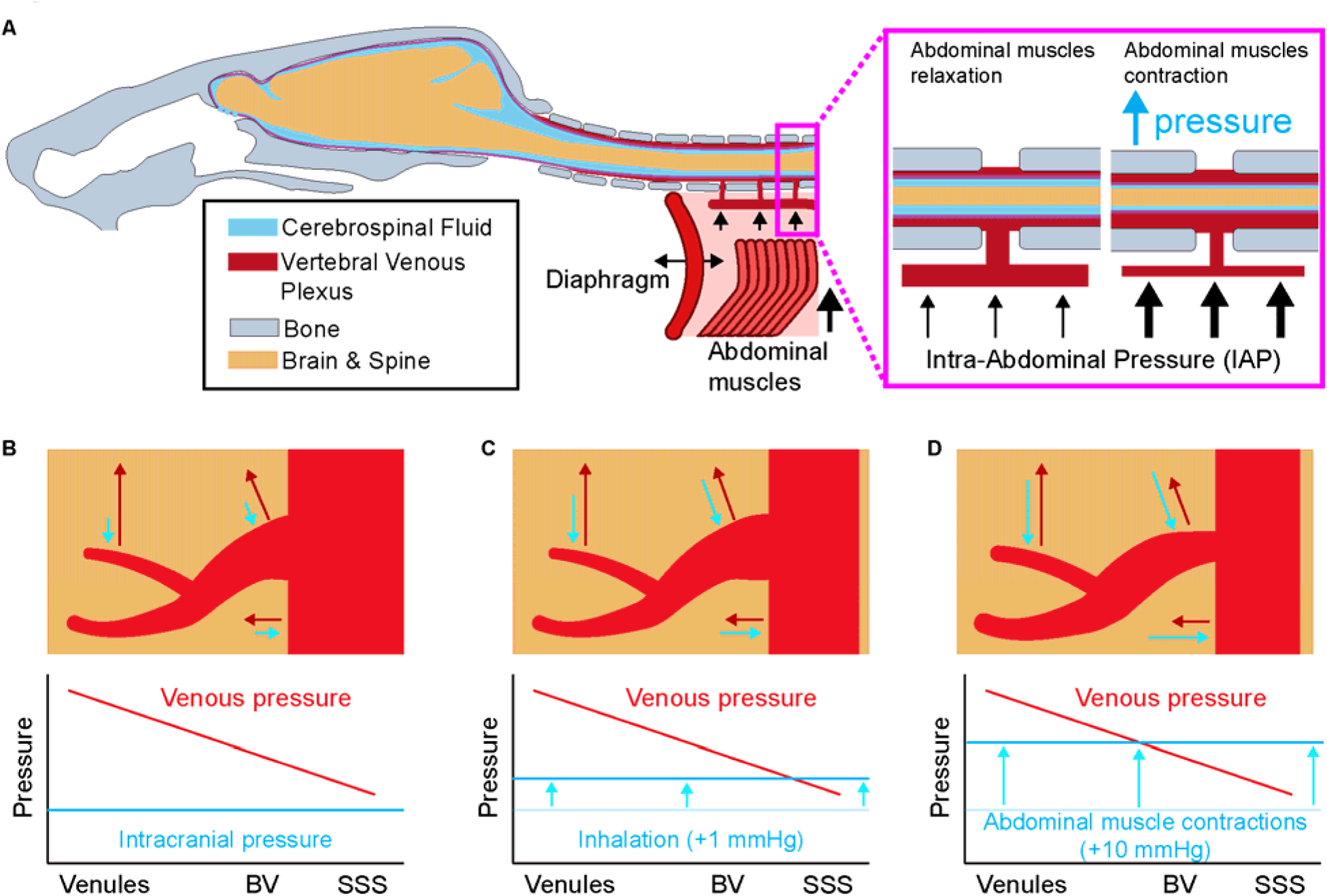
Elevated intracranial pressure can drive differential responses in venous diameter. (**A**) Schematic of the hypothesis that increases in intrabdominal pressure (due to abdominal muscle contraction) forces blood from the caudal vena cava to the vertebral venous plexus within the vertebral column. The increased blood volume in an enclosed space applies pressure to the dural sac, forcing the cranially-directed cerebral spinal fluid flow that increases intracranial pressure. (**B**) Intraluminal pressure of veins decreases as a function of location in the venous tree, with the largest veins having the lowest pressures. ICP on all these veins is approximately the same. (**C**) Respiration induces small variations of intracranial pressure, which will surpass the intraluminal pressure of superior sagittal sinus, leading to its compression. (**D**) During locomotion, sensory stimulation or other times abdominal muscles contract, ICP is elevated above the intraluminal pressures of the bridging veins and superior sagittal sinus, leading to their compression.

Previous work has observed rapid contractions of bridging veins (25, 55) and dural venous sinus (38, 56), though their origin was not understood. There have also been reports of stimulation-induced small, rapid blood volume decreases whose origin is mysterious (57), and of artifactually increased fluorescence just prior to locomotion onset in mouse two-photon imaging (37) that likely have contributions from the venous constriction we describe. Our work extends on previous work showing the brain is mechanically coupled to the abdominal cavity (23, 58, 59), and shows that the mechanical coupling to the abdomen can control blood flow in the brain with rapid dynamics.

Besides the increase in blood flow elicited by the compression of these veins, there may be other possible roles for this constriction. Firstly, the contractions of these veins could help compensate for ICP increases by providing compliance in the cranial cavity (60). Secondly, the SSS is a dural sinus, contraction of the SSS would apply tension to the mechanosensitive dura (61), which may account for some of the movement or deformation in the dura seen during locomotion in mice (62). Thirdly, the SSS is innervated by trigeminal and cervical spinal nerves (63) and is pain sensitive (64), so changes in the mechanical dynamics of the SSS could also play a role in migraine and headache (61). Fourthly, the contractions of these veins could also help move cerebrospinal fluid, which is known to be moved by dilations or contractions in arteries (65–67). The venous contractions may also help with the pumping action of the lymphatic vessels seen near the superior sagittal sinus (68–71). Finally, they may serve a mechanical signaling role. Mechanical coupling between the abdomen and brain is especially interesting considering the functional mechanosensitive channels in central nervous system neurons (72) and glia (73), and the rapid contractions of these veins could also activate mechanosensitive channels in the brain, acting as a direct interoceptive signaling pathway to the brain. These pathways may underlie cardiac and respiratory modulation of neural activity (74, 75).

While compression (stenosis) of veins by elevated intracranial pressure has been noted under pathological conditions (76–78), our work shows that this compression may occur naturally, albeit transiently, in the behaving brain. However, our results could have implications for brain pathologies as well. Pathologies like obesity elevate intra-abdominal pressure (79) and ICP (80), which could disrupt the normal flow of blood back and forth between the abdominal cavity and spinal canal. Elevated blood pressure may attenuate or block these venous contractions, potentially playing a role in the adverse effects of hypertension on brain health. From the clinical perspective, changes in the normal dynamics of SSS/BV constrictions could be used as non-invasive indicators of intracranial pressure (81–83). In summary, this work supports a model in which the brain and cerebral vasculature is mechanistically coupled with the viscera, providing a dynamic signaling pathway that can rapidly communicate information about the body state to the brain.

## Materials and Methods

### Animals

All experimental procedures were performed in accordance with the National Institute of Health guidelines and the Institution of Animal Care and Use Committee of the Pennsylvania State University (protocol #201042827). A total of 24 mice, including 18 (2 female) C57BL/6 mice (Jackson Laboratory, #000664), and 6 (all male) C57BL/6-Tg(CAG-EGFP)131Osb/LeySop mice (CAG-EGFP, Jackson Laboratory, #006567), were used. Recordings of brain hemodynamics response to locomotion were made from 7 C57BL/6 mice (2 female) using widefield imaging and laser Doppler flowmetry. In a subset of these mice (n = 4, all male), brain hemodynamics response to whisker stimulation, during sleep stage transition and to passive abdominal compression were made using widefield imaging and laser Doppler flowmetry. In a subset of these mice, recordings of vessel diameters using two-photon laser scanning microscopy were conducted in 5 mice (1 female). Miniscope measurements of venous diameters in free behaving mice were conducted in 1 male C57BL/6 mouse. Recordings of intracranial pressure were conducted in 10 male C57BL/6 mice. Recordings of fluorescence activity were conducted in 6 male CAG-EGFP mice. Mice were housed on a 12-hour light/dark cycle in isolated cages with access to food and water ad libitum. All experiments were conducted during the light cycle.

### Window, and electrode implantation procedures

Mice were 2 to 14 months old (20-35 g) at the time of surgery. All surgical procedures were performed under isoflurane anesthesia (induced by 5% isoflurane in oxygen and maintained at 1.5-2% during surgery). Body temperature was maintained by a thermal pad (Harvard Apparatus, Holliston, MA) set at 37 °C. A custom-made titanium head bolt was glued to the skull caudal from lambda using clear cyanoacrylate cement (catalog #32002, Vibra-Tite, Troy, MI) and black dental acrylic resin (catalog #1530, Lang Dental Manufacturing Co., Wheeling, IL) for head-fixation. To reduce motion artifacts, two self-tapping 3/32” #000 screws (J.I. Morris, Southbridge, MA) were placed into the skull over the olfactory bulb using cyanoacrylate cement and black dental acrylic resin. For brain hemodynamics measurements using optical imaging methods (widefield intrinsic optical signal imaging and two-photon laser scanning microscopy), a polished and reinforced thin skull (PoRTS) window was made covering both hemispheres as described previously (25, 27, 36, 51, 52, 84–86). For brain hemodynamics measurements using miniscope (nVue, Inscopix, Mountain View, CA), a PoRTS window was made covering the right hemisphere, and a baseplate was implanted for miniscope camera attachment. To measure neck muscle activity, the skin above the neck was resected and a pair of PFA-coated 7-strand stainless-steel wires (#793200, A-M Systems, Sequim, WA) were inserted into each nuchal muscle for electromyography (EMG) recording. To measure abdominal muscle activity, an incision 1 cm long was made in the skin below the ribcage to expose the oblique abdominal muscle. A small guide tube was then inserted into this incision and tunneled subcutaneously until it reached the open scalp. Two coated stainless steel electrode wires (#790500, A-M Systems, Sequim, WA) were inserted through the tube until the ends were exposed though both incisions, allowing the tube to be removed while the wires remained embedded under the skin. Two gold header pins (#0145-0-15-15-30-27-04-0, Mill-Max Manufacturing Corporation, Oyster Bay, NY) were adhered to the head bar with cyanoacrylate glue and the exposed wires between the header and neck incision were covered with silicone to prevent damage. Each wire exiting the abdominal incision was stripped of a section of coating and threaded through the muscle approximately 2 mm parallel from each other to allow for a bipolar abdominal EMG recording. A biocompatible silicone adhesive (KWIK-SIL, World Precision Instruments, Sarasota, FL) was used to cover the entry and exit of the muscle by the wires for implantation stability. The incision was then closed with a series of silk sutures (#18020-50, Fine Science Tools, Foster City, CA) and Vetbond (#1469, 3M, Livonia, MI). Following the surgery, mice were then returned to their home cage for recovery for at least one week and then started habituation on experimental apparatus.

### Habituation

Animals were gradually acclimated to head-fixation on a spherical treadmill (21, 25, 52, 86, 87) with one degree of freedom or a clear plastic tube (36, 47) (https://github.com/DrewLab/Mouse-Head-Fixation) over at least three habituation sessions. The spherical treadmill was covered with nonabrasive anti-slip tape (McMaster-Carr, Aurora, OH) and attached to an optical rotary encoder (#E7PD-720-118, US Digital, Vancouver, WA) to monitor locomotion. Mice were monitored by video for any signs of stress during habituation. Habituation was done over the course of one week, with the duration increasing from 5 min to 45 min. Mice that received whisker stimulation were acclimatized to head-fixation for 15–30 min during the first session. In subsequent sessions, they began to receive air puffs directed at the whiskers and were head-fixed for longer durations (> 60 minutes). For recordings using miniscope in free behaving mice, the animal was habituated to being handled and tethered to the miniscope in an open field chamber for three daily 30 minutes periods. In all cases, the mice exhibited normal behaviors such as exploratory whisking and occasional grooming after being head-fixed. Heart rate fluctuations were detectable in the intrinsic optical signal (25, 51) and varied between 7 and 13 Hz for all mice after habituation, which is comparable to the mean heart rate (∼12 Hz) recorded telemetrically from mice in their home cage (88).

### Histology

At the conclusion of the experiment, mice were deeply anesthetized with 5% isoflurane, transcardially perfused with heparinized saline, and then fixed with 4% paraformaldehyde. The brains were extracted and sunk in a 4% paraformaldehyde with 30% sucrose solution. The flattened cortices were sectioned tangentially (60 µm/section) using a freezing microtome and stained for the presence of cytochrome-oxidase (36, 89–91). The anatomical locations of the vasculature were then reconstructed using a combination of vascular images taken during surgery and the stained brain slices using Adobe Illustrator (Adobe Systems, San Jose, CA).

### Physiological measurements

Data from all experiments (except two photon laser scanning microscopy and miniscope imaging) were collected using custom software written in LabVIEW (version 2018, National Instruments, Austin, TX)(https://github.com/DrewLab/LabVIEW-DAQ).

#### Behavioral measurement

The treadmill movements were used to quantify the locomotion events of the mouse. The animal was also monitored using a camera (BFLY-PGE-23S6M-C PoE Mono, FLIR, Wilsonville, OR) during data acquisition as an additional behavioral measurement. Animal motion inside the tube was measured using a pressure sensor (Flexiforce A201, Tekscan, Boston, MA). The right vibrissae were diffusely illuminated from below by a 625 nm light (#66–833, Edmund Optics, Barrington, NJ), and imaged (30 x 350 pixels) using a camera (Basler ace acA640-120gm, Edmund Optics, Barrington, NJ) with a 18 mm DG Series FFL lens (#54–857, Edmund Optics, Barrington, NJ) at a nominal rate of 150 frames/second. The image was narrow enough to only show the whiskers as dark lines on a bright background, with the average whisker angle being estimated using the Radon transform (92). For both whisker acceleration and pressure sensor data, a threshold was manually set to establish when the animal behaved.

#### Widefield intrinsic optical signal imaging and fluorescence imaging

We mapped the spatiotemporal dynamics of oxyhemoglobin and deoxyhemoglobin concentrations using their oxygen-dependent optical absorption spectra (41). For imaging experiments using C57BL6 mice, reflectance images were collected during periods of green LED light illumination at 530 nm (equally absorbed by oxygenated and deoxygenated hemoglobin, M530L3, Thorlabs, Newton, NJ) or blue LED light illumination at 470 nm (absorbed more by oxygenated than deoxygenated hemoglobin, M470L3, Thorlabs, Newton, NJ). For these experiments, a CCD camera (Dalsa 1M60, Waterloo, ON) was operated at 60 Hz (equivalent two-color frame rate in each channel is 30 Hz) with 8.3 ms exposure and 4X4 binning (256 X 256 pixels), mounted with a VZM300i optical zoom lens (Edmund Optics, Barrington, NJ).

For experiments using CAG-EGFP mice, simultaneous widefield fluorescence and cortical hemodynamics signals were acquired by using a camera configured to acquire images in synchrony with three LEDs . Reflectance images were acquired during periods of green LED light illumination at 530 nm (M530L3, Thorlabs, Newton, NJ) or red LED light illumination at 630 nm (M625L4, Thorlabs, Newton, NJ), whereas a 500-nm long-pass filter (FELH0500, Thorlabs, Newton, NJ) mounted in front of the camera enabled imaging of GCaMP fluorescence during periods of illumination with blue LED light with a wavelength around 470 nm (M470L3, Thorlabs, Newton, NJ). For these experiments, a CCD camera (Dalsa 1M60, Waterloo, ON) was operated at 30 Hz (equivalent three-color frame rate in each channel is 10 Hz) with 10 ms exposure and 4X4 binning (256 X 256 pixels), mounted with a VZM300i optical zoom lens (Edmund Optics, Barrington, NJ).

#### Two-photon laser scanning microscopy imaging

Mice were briefly anesthetized with isoflurane (5% in oxygen) and retro-orbitally injected with 50 µL 5% (weight/volume in saline) fluorescein-conjugated dextran (70 kDa, Sigma-Aldrich, St. Louis, MO), and then fixed on a spherical treadmill. Imaging was done on a Sutter Movable Objective Microscope (Novato, CA) with a 16X, 0.8 NA water dipping objective (16XLWD-PF, Nikon, Tokyo, Japan). A MaiTai HP (Spectra-Physics, Santa Clara, CA) laser tuned to 800 nm was used for fluorophore excitation. All imaging with the water-immersion lens was done with room temperature distilled water. All the 2PLSM measurements were started at least 20 minutes after isoflurane exposure to avoid the disruption of physiological signals due to anesthetics (87, 93). For navigational purposes, widefield images were collected to generate vascular maps of brain pial vascular maps of the entire PoRTS window. We performed three different measurements using 2PLSM. To measure blood vessel diameter responses to locomotion, individual arteries and veins were imaged at nominal frame rate of 3 Hz for 5 minutes using 10-15 mW of power exiting the objective. Diameter of pial and dural vessels were calculated using the full width at half maximum (4).

#### Imaging brain hemodynamics in freely behaving mice using miniscope

Mice were briefly anesthetized with isoflurane (5% in oxygen) and retro-orbitally injected with 50 µL 5% (weight/volume in saline) rhodamine-conjugated dextran (70 kDa, Sigma-Aldrich, St. Louis, MO). The miniscope (Inscopix, Mountain View, CA) is then attached to the baseplate. Mice were then placed in the corner of a 50×50×20 cm open field arena and allowed to explore for 20 minutes. Behavior data and brain hemodynamics data were recorded using nVue system (Inscopix, Mountain View, CA) at 30 frames/second.

#### Electromyography (EMG)

EMG were recorded as the voltage difference between two stainless steel wires placed either at the neck muscle or external oblique of the abdominal muscle using an amplifier and an instrumentation amplifier. The acquired signals were amplified (gain = 1000, bandwidth 0.1-10k Hz, DAM80, World Precision Instruments, Sarasota, FL) and then digitized at 20 kHz (PCIe-6343, National Instruments, Austin, TX).

#### Respiration measurement using thermocouple

We conducted respiration recordings (25) during widefield intrinsic optical signal imaging experiments in a subset of mice (n = 4). Measurements of breathing were taken using 40-gauge K-type thermocouples (TC-TT-K-40-36, Omega Engineering, Norwalk, CT) placed near the mouse’s nose (∼ 1 mm), with care taken to not contact the whiskers. Data were amplified 2000x, filtered below 30 Hz (Model 440, Brownlee Precision, Santa Clara, CA), and sampled at 20 kHz (PCIe-6343, National Instruments, Austin, TX). Downward and upward deflections in respiration recordings correspond to inspiratory and expiratory phases of the respiratory cycle, respectively. We identified the time of each expiratory peak in the entire record as the zero-crossing point of the first derivative of the thermocouple signal(25).

#### Cerebral blood flow measurement using laser Doppler flowmetry

For cerebral blood flow (CBF) measurements, a laser Doppler flowmetry optical probe (OxyFlo, Oxford Optronix, Adderbury, UK) was placed at 45-degree angle to the window plane on the left hemisphere, avoiding major vessels on the brain surface (27).

#### Intracranial pressure (ICP) measurements

ICP data was taken from previous publications (21, 22). All surgical procedures were under isoflurane anesthesia (5% for induction and 2% for maintenance). A titanium head-bar was attached to the skull with cyanoacrylate glue and dental cement and the skull was covered with a thin layer of cyanoacrylate glue. After two days of recovery, the animal was habituated to head fixation on a spherical treadmill for one day (for three 30-minute sessions). On the day of the ICP experiment (one day after the habituation), the mouse was anesthetized with isoflurane and a small craniotomy (∼1 mm diameter) was made in the somatosensory cortex. A pressure measuring catheter (SPR-1000, Millar, Pearland, TX) was inserted into the cortex (−1.0 mm dorsal, -1 mm ventral from bregma), and a tight seal was made using Kwik-Cast low toxicity silicone sealant (World Precision Instruments, Sarasota, FL). This surgical procedure took approximately 10 minutes. The animal was allowed to wake from anesthesia and to freely locomote on the spherical treadmill for two hours, during which both intracranial pressure and locomotion were recorded simultaneously at 1 kHz (USB-6003, National Instruments, Austin, TX). To minimize any residual effect of anesthesia on ICP (21), we only analyzed data collected more than 1 hour after the cessation of anesthesia.

#### Whisker stimulation

Mice were stimulated with brief (0.1 s), randomized gentle puffs (10 psi) of air via an air regulator (Wilkerson R03-02-000, Grainger, Lake Forest, IL) to either the left vibrissae, right vibrissae, or an auditory control (with 1:1:1 ratio) every 30 s. The stimuli were directed to the distal ends of the whiskers, parallel to the face to avoid stimulating other parts of the body/face. Each stimulus was controlled with a solenoid actuator valve (2V025, Sizto Tech Corporation, Palo Alto, CA).

#### Abdominal pressure application

A custom-made pneumatically-inflatable belt (23) was fabricated to directly apply pressure to the abdomen of mice. It consisted of three plastic bladders that were fully wrapped around the abdomen of mice. The belt was inflated with 10 psi of pressure to apply a steady squeeze for 1 seconds with 45 seconds of rest between squeezes to allow for a return to baseline. The abdominal compression was oriented in such a way that no compression or tension was imparted to the spine longitudinally, as this could affect the results by pushing or pulling on the spine itself. Mice were observed with a behavioral video camera during imaging to check for potential compression-induced body positional changes.

### Data analysis

Data from all experiments (except the ones using Inscopix miniscope) were collected using custom software written in MATLAB (R2020b, Mathworks, Natick, MA).

#### Movement quantification

Locomotion events (21, 25, 36, 52) from the spherical treadmill were identified by first applying a low-pass filter (10 Hz, 5^th^ order Butterworth) to the velocity signal from the optical rotary encoder, and then comparing the absolute value of acceleration (first derivative of the velocity signal) to a threshold of 3 cm/s^2^. Periods of locomotion were categorized based on the binarized detection of the treadmill acceleration:

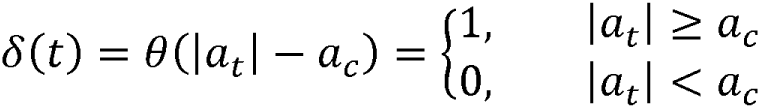

where a_t_ is the acceleration at time t, and a_C_ is the treadmill acceleration threshold. Movement data from the pressure sensor was digitally lowpass filtered (<20 Hz) using a second-order Butterworth filter and then resampled down to 30 Hz (MATLAB function(s): butter, zp2sos, filtfilt, resample). To identify movement events, the force sensor data was binarized acceleration by setting an empirically defined threshold.

#### Whisker motion quantification

Images of the mouse’s vibrissae were converted into a relative position (angle) by applying the Radon transform (MATLAB function(s): radon). The peaks of the sinogram corresponded to the position and the angle of the whiskers in the image (36). The average whisker angle was extracted as the angle of the sinogram with the largest variance in the position dimension (92). Any dropped frames were filled with linear interpolation between the nearest valid points (MATLAB function(s): interp1). Whisker angle was lowpass filtered (<20 Hz) using a second-order Butterworth filter and then resampled down to 30 Hz (MATLAB function(s): butter, zp2sos, filtfilt, resample). To identify periods of whisking, whisker acceleration was obtained from the second derivative of the position and binarized with an empirically chosen acceleration threshold for a whisking event. Acceleration events that occurred within 0.1 s of each other were linked and considered as a single whisking bout.

#### Widefield intrinsic optical signal and fluorescence imaging processing

We mapped the spatiotemporal dynamics of oxyhemoglobin and deoxyhemoglobin concentrations using their oxygen-dependent optical absorption spectra (41). Reflectance data (either green and blue, or green and red when performing fluorescence imaging) were converted to changes in oxy- and deoxyhemoglobin concentrations using the modified Beer-Lambert law with Monte Carlo-derived wavelength-dependent path length factors (41). We used the cerebral oxygenation index (94) (i.e., HbO-HbR) to quantify the change in oxygenation, as calculating the percentage change requires knowledge of the concentration of hemoglobin on a pixel-by-pixel basis, which is not feasible given the wide heterogeneity in the density of the cortical vasculature (95, 96).

Interspersed fluorescence images of fluorescence signals were hemodynamic corrected and converted into ΔF/F_0_, where ΔF is the mean subtracted raw fluorescence signal and F_0_ is the mean of the entire time course. To correct hemodynamics contamination, we tested three widely used methods: (1) hemodynamic correction using only green reflectance (i.e., single-wavelength method)(43), (2) hemodynamic correction using estimated excitation and emission attenuation (i.e., Ex-Em method)(40, 41), and (3) hemodynamic correction using a spatially detailed regression-based method to estimate hemodynamics contamination (i.e., spatial model)(38). The spatial model is based on a linear form of the Beer-Lambert relationship but is fit at every pixel in the image and does not rely on the estimation of physical parameters.

To calculate venous diameter using widefield intrinsic optical imaging, vessel diameters were calculated from the normalized full width at half minimum of the image intensity along a line perpendicular to the vessel, average of results from 5 parallel lines were used for vessel diameter (25).

#### Two-photon laser scanning microscopy imaging processing

Individual stack frames from 2PLSM were corrected for x-y motion and aligned with a rigid registration algorithm (4, 8).

Imaging periods with excessive z-plane motion artifacts were excluded from analysis. A rectangular box was manually drawn around a straight, evenly-illuminated segment of the vessel and the pixel intensity was averaged along the long axis and used to calculate the vessel’s diameter from the full-width at half-maximum (4) (https://github.com/DrewLab/Surface-Vessel-FWHM-Diameter).

#### Miniscope data processing

The imaging data were processed and analyzed using the Inscopix Data Processing Software (Inscopix, Mountain View, CA).

#### Electromyography (EMG)

Electrical activity from the nuchal (neck) muscles or abdominal muscles was digitally bandpass filtered (300 Hz – 3 kHz) using a third-order Butterworth filter. The signal was then squared and convolved with a Gaussian kernel with a 0.5 s standard deviation, log transformed, and resampled down to 30 Hz (MATLAB function(s) butter, zp2sos, filtfilt, gausswin, log10, conv, resample). Folder increase was determined as Power/Power_baseline_.

#### Laser Doppler flow velocimetry (LDF)

Microvascular perfusion data was resampled down to 30 Hz and digitally lowpass filtered (<1 Hz) using a fourth-order Butterworth filter (MATLAB function(s) butter, zp2sos, filtfilt, resample).

#### Sleep scoring methodology

We use previously published methods (47) for sleep stage identification.

#### Spontaneous and evoked activity

To characterize spontaneous (non-locomotion-evoked) activity, we defined resting periods as periods started 4 seconds after the end of previous locomotion event and lasting more than 10 seconds. Locomotion-evoked events were defined as segments with at least 3 seconds of resting prior to the onset of locomotion and followed by at least 5 seconds of locomotion. Whisker stimulation evoked events were defined as segments the animal receiving air puff with no significant movements 3 seconds before and 5 seconds after stimulation.

#### Cross-correlation analysis

Cross-correlation analysis was performed between simultaneously recorded neural/respiration and hemodynamics signals (ΔHbT, venous diameter) to quantify the relationship between fluctuations. For spontaneous correlations, only periods of rest lasting more than 30 seconds were used, with a four-second buffer at the end any locomotion event. We also calculated the correlations using all the data including periods with locomotion. The temporal cross-correlation between EMG power/respiration/locomotion speed and brain hemodynamics signals was calculated (xcorr, MATLAB). Positive delays denote the EMG/respiration/locomotion signal lagging the brain hemodynamics signal. Statistical significance of the correlation was computed using bootstrap resampling from 1000 reshuffled trials.

### Statistical analysis

Statistical analysis was performed using MATLAB (R2020b, Mathworks). All summary data were reported as the mean ± standard deviation (SD) unless stated otherwise. To test whether the brain hemodynamics are increasing or decreasing, Wilcoxson rank sum test (MATLAB: ranksum) was used to test against zero. To compare the performance of the methods used to remove the hemodynamics contamination from fluorescent signals, paired t-test (MATLAB: ttest) was used. Significance was accepted at *p* < 0.05.

## Acknowledgments

This work is supported by National Institute of Health grants R01NS078168 and U19NS128613 to PJD, and American Heart Association Career Development Grant 935961 to QZ.

## Author Contributions

Q.Z. and P.J.D. designed the project. Q.Z. performed experiments and analyzed data using widefield imaging, two photon laser scanning microscopy, miniscope, and laser Doppler flowmetry. C.S.G. helped with EMG implantation. N.F. designed the pneumatic cuff. F.S. performed sleep experiments. K.L.T. performed fluorescence signal experiments. P.J.D. supervised experiments, data analysis, and preparation of the manuscript. Q.Z. and P.J.D. wrote the manuscript with input from all authors.

## Supplementary figure captions

**Supplementary Figure 1.**
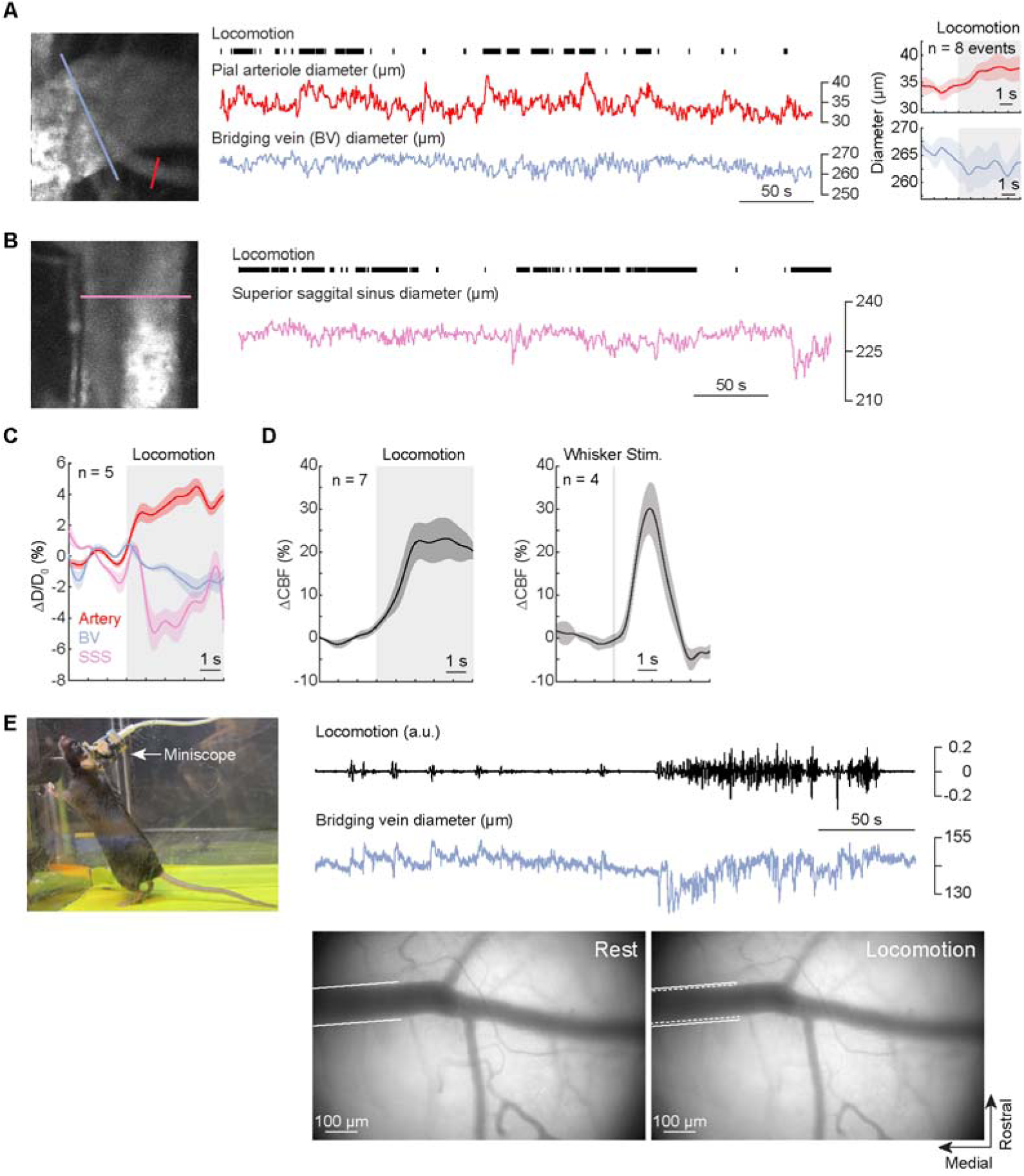
Two-photon and Miniscope imaging of venous responses to voluntary locomotion. (**A**) Left, example bridging vein (purple) and pial artery (magenta) dynamics visualized with two-photon microscopy. Black tick marks denote locomotion. Right, locomotion-triggered responses of the artery and bridging vein. (**B**) Two-photon imaging of the superior sagittal sinus. (**C**) Group (n = 5 mice) average of locomotion evoked responses in pial artery, bridging vein and superior sagittal sinus. There was a rapid constriction in superior sagittal sinus (SSS, -0.37 ± 0.42%, Wilcoxson rank sum test, p = 0.0635) or bridging veins (BV, -0.76 ± 0.84%, Wilcoxson rank sum test, p = 0.0084) in response to locomotion onset (0-1 second after locomotion onset), while pial arteries in the FL/HL dilates (2.00 ± 0.66%, Wilcoxson rank sum test, p = 0.09). In response to constant locomotion (2-5 seconds after locomotion onset), the artery maintained dilation status (3.76 ± 0.95%, Wilcoxson rank sum test, p = 0.0079), while the constricted SSS and BV remained constricted (SSS: -3.04 ± 1.26%, Wilcoxson rank sum test, p = 0.0079; BV, -1.57 ± 0.90%, Wilcoxson rank sum test, p = 0.0159). (**D**) Cerebral blood flow increases during locomotion and contralateral whisker stimulation. Significant increases of ΔCBF were observed in the parenchyma in response to constant locomotion (2-5 seconds after locomotion onset, 19.35 ± 9.44%, Wilcoxson rank sum test, p = 0.0079) and contralateral whisker stimulation (19.05 ± 8.79%, Wilcoxson rank sum test, p = 0.0286). (**E**) Example miniscope imaging in a freely moving mouse showing bridging vein constriction in response to voluntary locomotion.

**Supplementary Figure 2.**
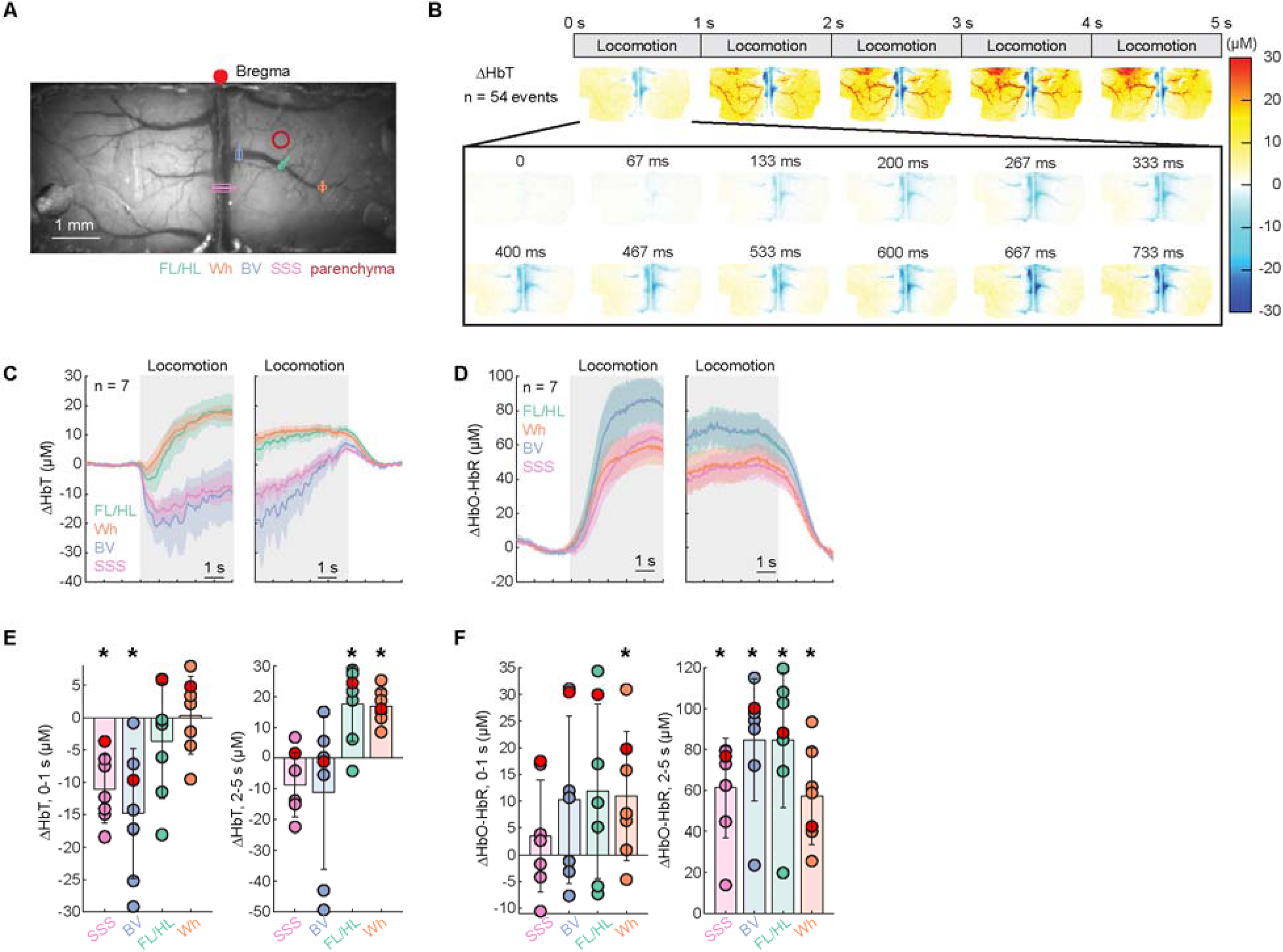
Venous responses to voluntary locomotion quantified by hemoglobin signals. (**A**) Image of the cerebral vasculature under 530 nm illumination through a thin-skull window spanning the parietal cortices of both hemispheres. Colored lines denote locations of vessel diameter measurements shown in subsequent figures. (**B**) Locomotion-triggered spatial pattern average of ΔHbT aligned to locomotion onset (n = 54 events in 1 mouse). Note visible changes within ∼100 miliseconds. (**C**) Average of locomotion onset and offset triggered response of ΔHbT (n = 7 mice) in veins at different locations. (**D**) As in (**C**) but for ΔHbO-HbR (n = 7 mice). (**E**) Average total hemoglobin change (ΔHbT) during the initial phase of locomotion (0-1 second after locomotion onset, left) and sustained locomotion (2-5 seconds after locomotion onset, right) in veins at different locations. In response to voluntary locomotion onset, we observed a fast decrease of ΔHbT in superior sagittal sinus (SSS, -11.07 ± 5.33 µM, Wilcoxson rank sum test, p = 0.0003) and bridging veins (BV, -14.80 ± 10.02 µM, Wilcoxson rank sum test, p = 0.0003), while ΔHbT of pial veins in the FL/HL (−3.60 ± 8.92 µM, Wilcoxson rank sum test, p = 0.09) and Wh (0.33 ± 6.01 µM, Wilcoxson rank sum test, p = 0.7164) did not change significantly. In response to constant locomotion, the initially decreased ΔHbT returned to baseline level for both the SSS (−8.82 ± 10.45 µM, Wilcoxson rank sum test, p = 0.09) and the BV (−11.18 ± 24.87 µM, Wilcoxson rank sum test, p = 0.3445), while ΔHbT of pial veins in the FL/HL (17.60 ± 12.21 µM, Wilcoxson rank sum test, p = 0.0169) and Wh (16.87 ± 5.51 µM, Wilcoxson rank sum test, p = 0.0006) increase significantly. (**F**) As in (**E**) but for ΔHbO-HbR. In response to voluntary locomotion onset, no significant change of ΔHbO-HbR was observed in superior sagittal sinus (SSS, 3.48 ± 10.50 µM, Wilcoxson rank sum test, p = 0.7164), bridging veins (BV, 10.28 ± 15.68 µM, Wilcoxson rank sum test, p = 0.7164), and pial veins in the FL/HL (11.84 ± 16.30 µM, Wilcoxson rank sum test, p = 0.9327) and Wh (10.97 ± 12.08 µM, Wilcoxson rank sum test, p = 0.9980). During locomotion, significant increases of ΔHbO-HbR were observed in superior sagittal sinus (SSS, 61.33 ± 24.34 µM, Wilcoxson rank sum test, p = 0.0006), bridging veins (BV, 84.79 ± 29.85 µM, Wilcoxson rank sum test, p = 0.0006), and pial veins in the FL/HL (84.66 ± 33.03 µM, Wilcoxson rank sum test, p = 0.0006) and Wh (57.28 ± 23.55 µM, Wilcoxson rank sum test, p = 0.0006).

**Supplementary Figure 3.**
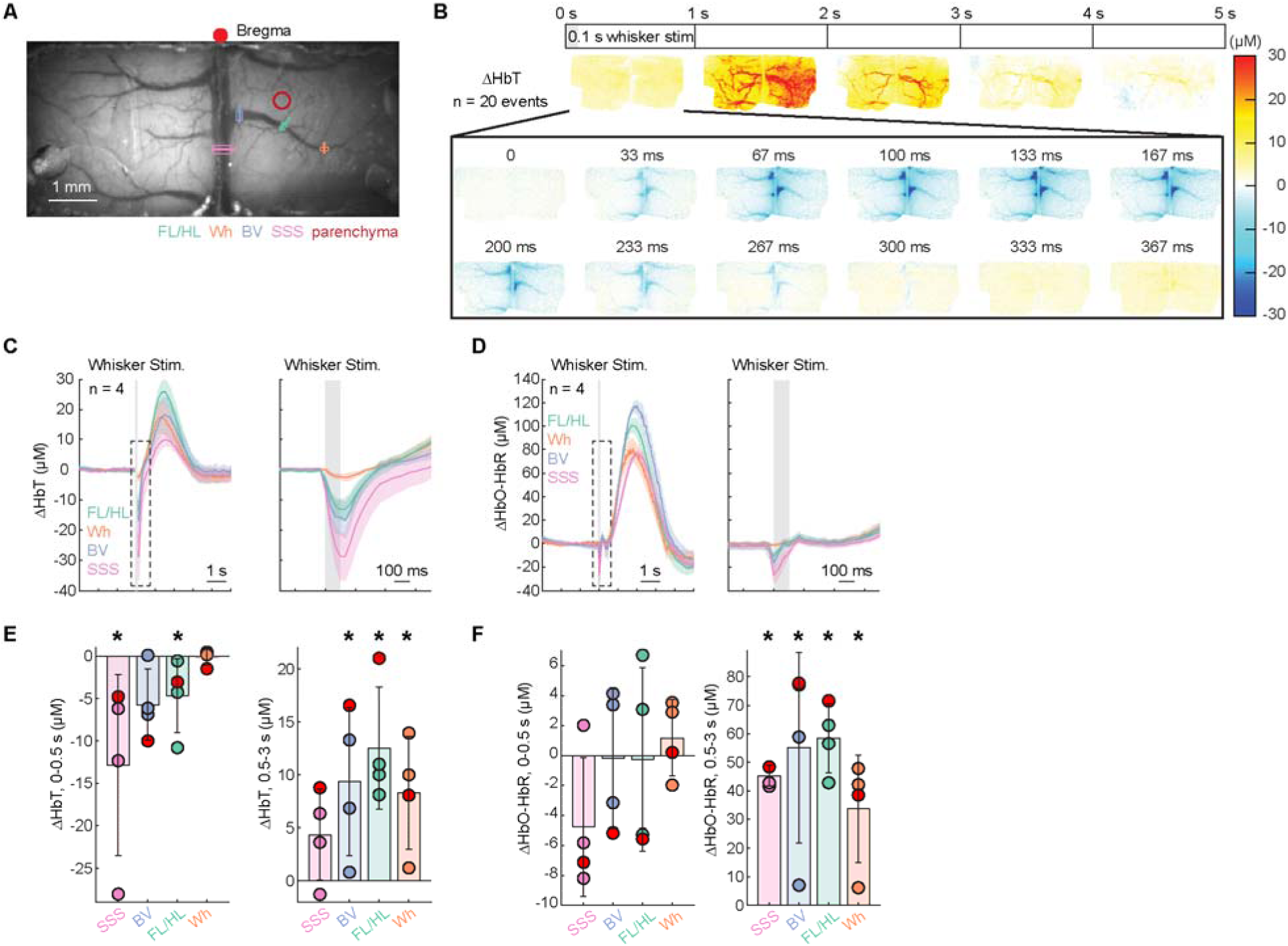
Brain hemodynamic responses to different whisker stimulation patterns. (**A**) Image of the cerebral vasculature under 530 nm illumination through a thin-skull window spanning the parietal cortices of both hemispheres. Colored lines denote locations of vessel diameter measurements shown in subsequent figures. (**B**) Averaged spatial distribution of ΔHbT in response to contralateral whisker stimulation (n = 20 events in 1 mouse). (**C**) Group average of contralateral whisker stimulation triggered response of ΔHbT (n = 4 mice) in veins at different locations. Inlet showing a zoom-in view of ΔHbT responses immediately before (330 ms) and after (660 ms) whisker stimulation. (**D**) As in (**C**) but for ΔHbO-HbR (n = 4 mice). (**E**) Average total hemoglobin change (ΔHbT) during the initial phase of contralateral whisker stimulation (0-0.5 second after contralateral whisker stimulation onset, left) and 0.5-3 seconds after whisker stimulation (right) in veins at different locations. During the initial phase after the whisker stimulation, we observed a fast decrease of ΔHbT in superior sagittal sinus (SSS, - 12.83 ± 10.66 µM, Wilcoxson rank sum test, p = 0.0143) and pial veins in the FL/HL (−4.66 ± 4.37 µM, Wilcoxson rank sum test, p = 0.0143), while the ΔHbT of bridging veins (BV, -5.71 ± 4.22 µM, Wilcoxson rank sum test, p = 0.1571) and pial veins in the Wh (−0.11 ± 0.92 µM, Wilcoxson rank sum test, p = 0.9429) did not change significantly. During later phase after the whisker stimulation, the ΔHbT returned to baseline level in SSS (4.38 ± 4.30 µM, Wilcoxson rank sum test, p = 0.1571), while ΔHbT of BV (9.38 ± 6.99 µM, Wilcoxson rank sum test, p = 0.0143), as well as ΔHbT of pial veins in the FL/HL (12.53 ± 5.78 µM, Wilcoxson rank sum test, p = 0.0143) and Wh (8.32 ± 5.31 µM, Wilcoxson rank sum test, p = 0.0143) increase significantly. (**F**) As in (**E**) but for ΔHbO-HbR. During the initial phase after whisker stimulation, no significant change of ΔHbO-HbR was observed in superior sagittal sinus (SSS, -4.77 ± 4.64 µM, Wilcoxson rank sum test, p = 0.1571), bridging veins (BV, -0.20 ± 4.66 µM, Wilcoxson rank sum test, p = 0.6), and pial veins in the FL/HL (−0.25 ± 6.13 µM, Wilcoxson rank sum test, p = 0.6) and Wh (1.17 ± 2.55 µM, Wilcoxson rank sum test, p = 0.9429). During later phase after whisker stimulation, significant increases of ΔHbO-HbR were observed in superior sagittal sinus (SSS, 45.26 ± 3.62 µM, Wilcoxson rank sum test, p = 0.0286), bridging veins (BV, 55.15 ± 33.25 µM, Wilcoxson rank sum test, p = 0.0286), and pial veins in the FL/HL (58.46 ± 12.10 µM, Wilcoxson rank sum test, p = 0.0286) and Wh (33.64 ± 18.73 µM, Wilcoxson rank sum test, p = 0.0286).

**Supplementary Figure 4.**
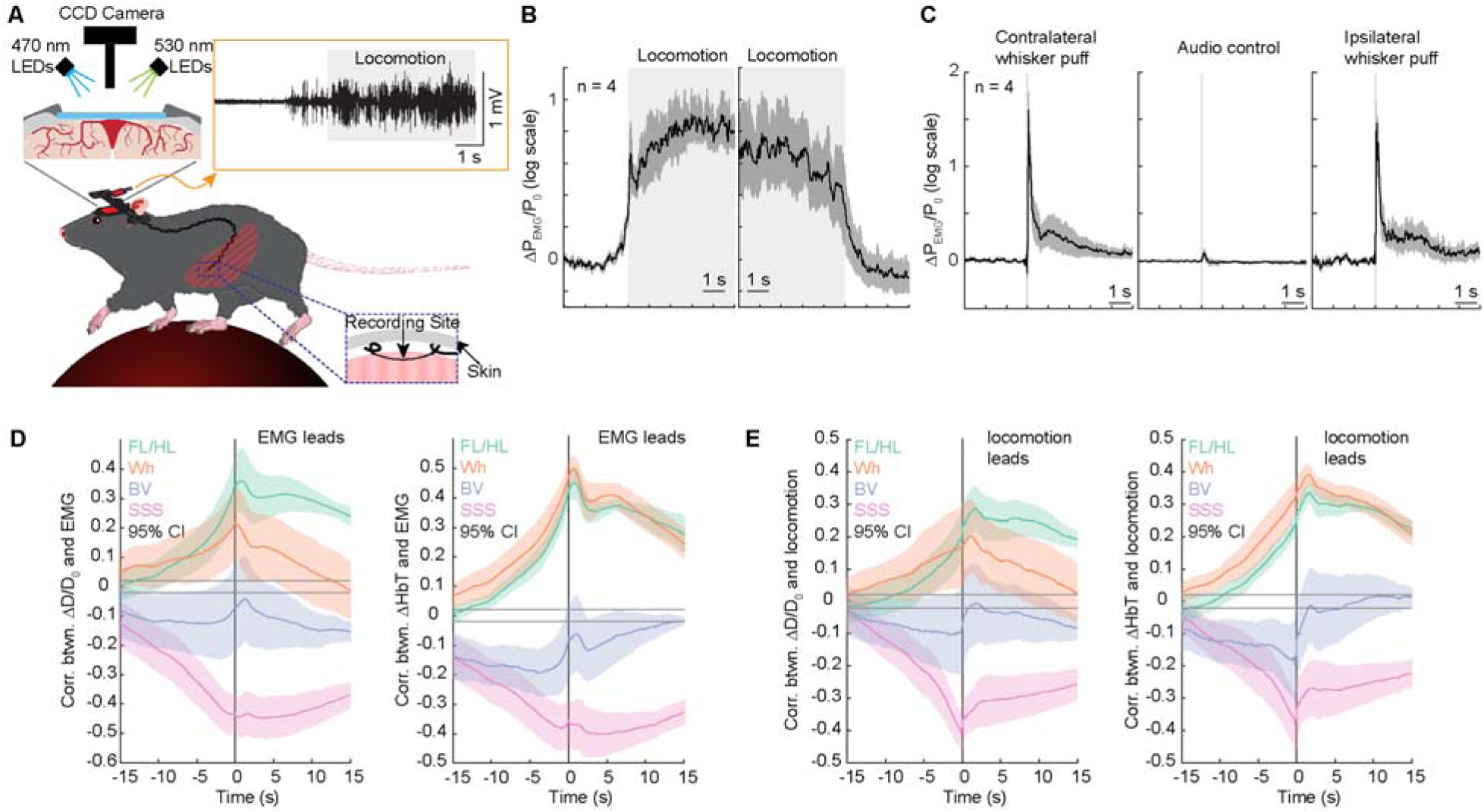
Abdominal muscle EMG precedes the venous constriction and locomotion. (**A**) Schematic of the experimental setup for simultaneous widefield IOS imaging and abdominal muscle EMG measurements of head fixed mice. Inset showing the EMG response during locomotion. (**B**) Locomotion onset (left) and offset (right) evoked responses of abdominal muscle EMG (n = 4 mice). Note EMG power increases prior to the onset of locomotion. (**C**) Whisker stimulation evoked responses of abdominal muscle EMG (n = 4 mice). (**D**) Cross-correlation between venous diameter change (left) or ΔHbT (right), and abdominal muscle EMG activity. An increase in abdominal muscle EMG activity leads to constrictions of SSS and BV, but dilations of pial veins. (**E**) As in (**D**) but for the cross-correlation between venous diameter change (left) or ΔHbT (right), and locomotion activity. An increase in locomotion speed occurs simultaneously as the constriction of SSS and BV, but leads to dilations of pial veins.

**Supplementary Figure 5.**
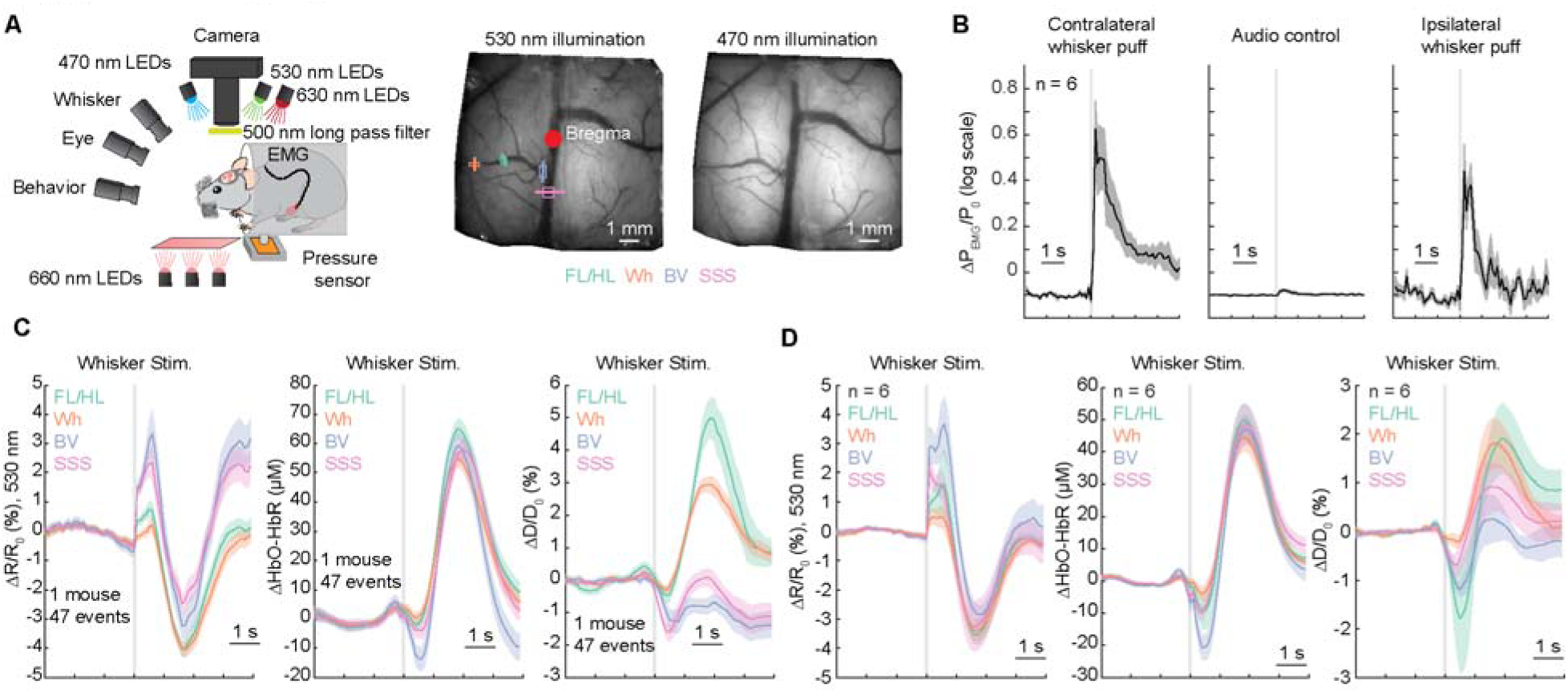
Brain hemodynamic responses to different whisker stimulation patterns while the mouse in the tube. (**A**) Schematic of the experimental setup for simultaneous widefield IOS imaging and abdominal muscle EMG measurements of head fixed mice. (**B**) Whisker stimulation evoked responses of abdominal muscle EMG (n = 6 mice). (**B**) Averaged response of cerebral blood volume (ΔR/R_0_ at 530 nm), brain tissue oxygenation (ΔHbO-HbR) and venous diameter (ΔD/D_0_) in response to whisker stimulation (n = 47 events in 1 mouse). (**C**) Group (n = 6 mice) average of whisker stimulation triggered response of of cerebral blood volume (ΔR/R_0_ at 530 nm), brain tissue oxygenation (ΔHbO-HbR) and venous diameter (ΔD/D_0_) in veins at different locations.

